# Mechanistic insights into RNA binding and RNA-regulated RIG-I ubiquitination by TRIM25

**DOI:** 10.1101/2020.05.04.070177

**Authors:** Kevin Haubrich, Sandra Augsten, Lucía Álvarez, Ina Huppertz, Bernd Simon, Kathryn Perez, Pawel Masiewicz, Mathilde Lethier, Katrin Rittinger, Frank Gabel, Matthias W. Hentze, Stephen Cusack, Janosch Hennig

## Abstract

TRIM25 is a ubiquitin E3 ligase active in innate immunity and cell fate decisions. Mounting evidence suggests that TRIM25′s E3 ligase activity is regulated by RNAs. However, while mutations affecting RNA binding have been described, neither the precise RNA binding site has been identified nor which domains are involved. Here, we present biophysical evidence for the presence of RNA binding sites on both TRIM25 PRY/SPRY and coiled-coil domains, and map the binding site on the PRY/SPRY with residue resolution. Cooperative RNA-binding of both domains enhances their otherwise transient interaction in solution and increases the E3 ligase activity of TRIM25. We also show that TRIM25 not only binds RNA in mammalian cells but that interfering with RNA binding has an effect on cellular RIG-I ubiquitination.

## INTRODUCTION

RIG-I signalling mediated by TRIM25 and related E3 ligases is one of the key steps in the host response against a broad spectrum of RNA viruses, many of which pose a significant hazard for public health and human well-being, such as Influenza, Dengue, Ebola and the novel coronaviruses (1,2). Many of these viruses have developed host-pathogen interactions to evade host immunity by interfering with TRIM25-mediated RIG-I ubiquitination (3–9). The recent outbreaks of emerging human pathogens such as Ebola in Western Africa in 2013-2016 or the ongoing SARS-CoV-2 pandemic, the steady rise in Dengue virus infections over the last 50 years as a consequence of human-made climate change, as well as the seasonal death toll caused by Influenza virus, which could be significantly increased in the event of a pandemic, highlight the importance of understanding these host-pathogen interactions as a potential target for fighting these diseases.

In this study we focus on the role of RNA binding of the E3 ligase TRIM25 in its physiological functions and its exploitation by viruses in host immunity evasion (10–13). TRIM25 is part of the tripartite motif (TRIM) family of ubiquitin ligases characterized by an N-terminal RING domain, followed by one or two B-box domains, a coiled-coil (CC) dimerization domain and a C-terminal region, which, depending on the subfamily, can feature various domains (Figure 1A). TRIM25 is a member of the PRY/SPRY domain subfamily and the first member of this subgroup for which RNA binding has been observed (7,14–16). TRIM25 has reported functions in innate immunity, morphogenesis and cell proliferation (17–21), but mechanistic understanding of its mode of action is scarce.

**Figure 1.**
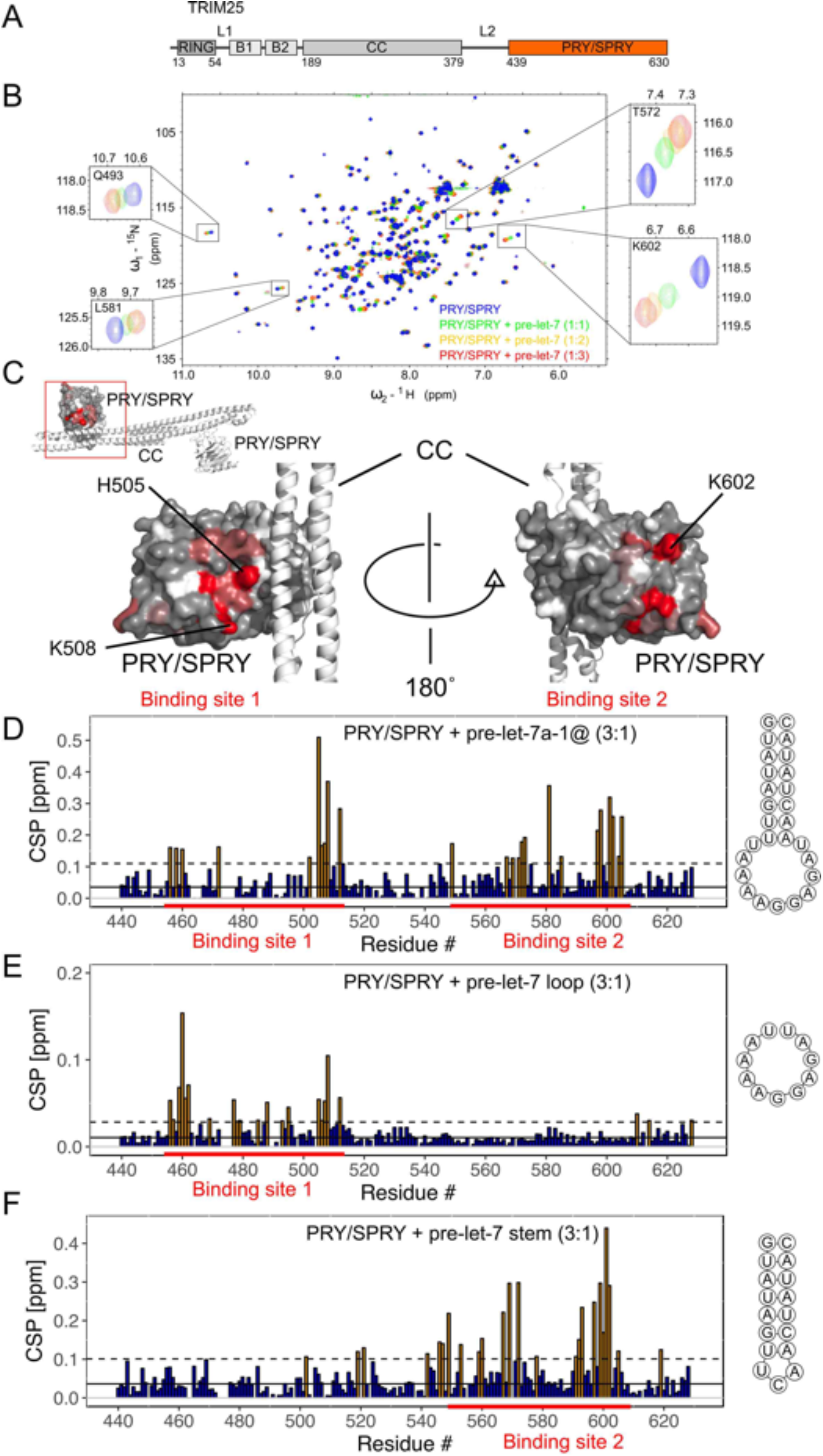
NMR titrations of TRIM25 PRY/SPRY domain and pre-let-7 RNA: **(A)** TRIM25 is a member of the tripartite motif family of E3 ligases characterized by a RING domain, two B-Boxes and a Coiled-coil. In addition, it has a C-terminal PRY/SPRY domain, which harbours RNA binding activity characterized in this figure **(B)** The ^1^H-^15^N-HSQC spectra of TRIM25 PRY/SPRY (130 μM) show strong chemical shift perturbations (CSPs) upon addition of pre-let-7 RNA up to a protein:RNA ratio of 1:3. **(C)** Mapping CSPs on the structure (PDB: 6FLM, coloured as a gradient from grey to red, unassigned residues in white), two strongly affected regions on the structure were identified. The strongest perturbations are located in binding site 1 close to the interface with the CC, while the second region is found on the opposing face of the domain. Comparison of the CSPs upon binding to full-length pre-let-7 **(D)** with those caused by the loop and stem region shows that the single-stranded loop region causes perturbations of binding site 1 (**E**), while the double-stranded stem affects almost exclusively binding site 2 **(F)**. Significantly affected residues are plotted in yellow and the binding sites in the sequence are indicated by red bars. Perturbation close to the C-terminus occur as the C-terminus folds back into the proximity of binding site 1. CSPs more than one standard deviation (dashed black line) above the average (continuous black line) in C, D, and E are coloured in orange.

The best characterized function of TRIM25 is its role in RIG-I signalling during antiviral response. Here, it ubiquitinates the RIG-I caspase activation and recruitment domains (CARDs), which are exposed upon recognition of 5′ triphosphate blunt-end double-stranded RNA by RIG-I’s helicase and C-terminal domain (17,22–26). Ubiquitination enhances oligomerization of the RIG-I CARDs, which activates the mitochondrial antiviral signaling protein MAVS, leading to interferon expression (27,28). The importance of this pathway in innate immunity is highlighted by the variety of mechanisms that viruses have evolved to inhibit TRIM25 activity and evade the host defense (3–8). TRIM25 has been shown to also ubiquitinate other targets, often RNA-binding proteins. In embryonic stem cells, TRIM25 is required for the binding of Lin28a to the precursor of let-7 (pre-let-7), which in turn recruits TUT4 to the RNA (29). TUT4, after activation through ubiquitination by TRIM25, poly-uridylates pre-let-7 and thereby marks it for degradation. Furthermore, TRIM25 ubiquitinates the zinc-finger antiviral protein (ZAP), a protein interacting with viral mRNA and facilitating exosome-mediated degradation (30). Various RNAs seem to act as co-factors for TRIM25’s E3 ubiquitin ligase activity, facilitating auto-ubiquitination and ubiquitination of RIG-I and ZAP (31–33). While most studies point towards an activating role of RNA on TRIM25 *in vitro*, there is some evidence that the Dengue virus uses its subgenomic RNA to inhibit TRIM25 and its role in interferon expression (4).

Despite clear evidence for the importance of RNA-binding for TRIM25 activity, the molecular mechanism of this regulation remains unexplained and structural data on the TRIM25-RNA interaction is lacking. The identity of the RNA-binding domain also remains controversial, with the CC domain originally being identified (14), while more recent data point towards the PRY/SPRY domain (31). In addition, a lysine-rich part of the linker connecting CC and PRY/SPRY domain was proposed to be involved in this interaction (32). Here, we resolve this controversy by showing that the CC and PRY/SPRY domains bind RNA cooperatively. We have previously shown that the PRY/SPRY and CC interact weakly in solution (34) and that this interaction is necessary for RIG-I ubiquitination. Binding to RNA enhances the CC-PRY/SPRY interaction, mechanistically explaining the observed activation of TRIM25 E3-ubiquitin ligase activity. We also show that TRIM25 binds preferably to stem-loops including a sequence-specificity towards A/G-rich sequences within the loop. Functional assays in mammalian cells confirm the importance of RNA binding interfaces on both domains, as RNA binding mutants lessen cellular RNA binding as well as RIG-I ubiquitination.

## MATERIAL AND METHODS

### Protein expression and purification

TRIM25 CC (aa 189-379), PRY/SPRY (aa 439-630) and CC-PRY/SPRY (aa 189-630) were cloned using restriction free cloning into pETM22 featuring a 3C-protease cleavable hexa-histidine and thioredoxin tag (His-Trx). An alternative N-terminally extended PRY/SPRY construct (aa 407-630) was cloned into pETM20 with a TEV-protease cleavable His-Trx tag. TRIM25 189-379 was expressed in BL21(DE3). TRIM25 439-630, 407-630 and 189-630 were co-expressed with chaperones KJE, ClpB and GroELS in *E. coli* BL21(DE3) (35). All proteins were induced at OD_600_ = 0.6 by 200 μM isopropyl-ß-D-1-thiogalactopyranoside (IPTG) and incubated at 18°C for 20h.

All proteins were purified by immobilized Nickel affinity chromatography (GE Histrap FF) in 50 mM Tris, pH 7.5, 300 mM NaCl and 0.2 mM TCEP and eluted with a gradient of imidazole (10-300 mM). In the case of co-expression with chaperones the column was washed with 50 mM Tris, pH 7.5, 350 mM KCl, 5 mM MgSO4 and 1 mM ATP prior to elution. The tag was removed by TEV-protease digestion overnight and an additional passage over the HisTrap column (189-379 and 407-630) or GE HiTrap SP HP cation exchange column (PRY/SPRY, residues 439-630 and CC-PRY/SPRY, residues 189-630). As a final step aggregates were removed by gel filtration on a GE Superdex S75 (CC, PRY/SPRY, extended PRY/SPRY) or S200 (CC-PRY/SPRY) in 20 mM MES, pH 6.5, 75 mM NaCl and 0.5 mM TCEP. For stable isotope labelling, proteins were expressed in M9 medium using ^15^NH_4_Cl (0.5 g/l) or ^15^NH_4_Cl/^13^C-Glucose (2 g/l) as sole nitrogen and carbon source.

Human RIG-I CARDs (aa 1-203 and 1-208) were cloned into pETM11 using the NcoI and KpnI sites, expressed in BL21(DE3) Rosetta 2 and induced with 250 μM IPTG overnight at 16 °C. After lysis the cleared lysate was treated with 1 M Urea in 25 mM Tris pH 7.5, 150 mM NaCl, 10% glycerol, 0.5 mM TCEP. The protein was purified using a Qiagen Nickel NTA superflow column in 25 mM Tris pH 7.5, 150 mM NaCl, 10% glycerol, 0.5 mM TCEP. The column was washed with 20 mM imidazole and 1 M salt and the protein eluted with 300 mM imidazole in the same buffer. The tag was removed by TEV-protease cleavage and an additional passage over the Ni-NTA column. Residual aggregates were removed in a final size exclusion chromatography step on a Superdex 75 column (GE healthcare) in a buffer containing 25 mM HEPES pH 7.5, 150 mM NaCl, 0.3 mM TCEP.

### RNA production

Pre-let-7a-1@2 (5′-GUA UAG UUU AAA AGG AGA UAA CUA UAC –3′), Lnczc3h7a-SL (5′-UUUUAUCUGAGUUGGAGGUGAAG-3′) and DENV-SL (5′-GCA GGU CGG AUU AAG CCA UAG UAC GGG AAA AAC UAU GCU ACC UG-3′) were *in vitro* transcribed from DNA oligos using T7 RNA polymerase. Reaction mixtures containing 2 μM forward and reverse primer, 40 mM Tris, pH 8.0, 0.2 mM MgCl2, 10 mM NTPs, 10 mM spermidine, 15 mM DTT, 0.01 % Triton X-100, 4 U/ml TIPP, 0.1 mg/ml T7 polymerase were incubated for 5 hours at 37°C and extracted by chloroform/phenol treatment. The aqueous phase was further purified by preparative gel electrophoresis via denaturing PAGE or HPLC using a Thermo DNA Pac PA100 22×250mm anion exchange column at 95° C. The purified RNA was dialyzed against 20 mM MES, pH 6.5, 75 mM NaCl and 0.5 mM TCEP and refolded before use by heating up at 95°C for 5 minutes and snap-cooling on ice. The shorter pre-let-7 loop (5′-UAA AAG GAG AU-3′), stem-fusion (5’-G UAU AGU U C AAC UAU AC-3′), Lnczc3h7a-SL loop (5′-UGAGUUGGA-3′) and 28-mer duplex RNA (5′-AUG GCU AGC UGG AGC CAC CCG CAG UUC G-3′) constructs were purchased from IBA and Biomers.

### Nuclear magnetic resonance spectroscopy

NMR spectra were acquired on Bruker Avance III spectrometers operating at magnetic field strengths corresponding to proton Larmor frequencies of 600, 700, and 800 MHz, equipped with a cryogenic triple resonance probe (600 and 800 MHz) and a room temperature triple resonance probe (700 MHz). Protein observed experiments were performed in 20 mM sodium phosphate, pH 6.5, 150 mM NaCl, 2 mM TCEP, 5% D_2_O and 0.02% sodium azide at 293 K. For RNA observed experiments samples were prepared in 20 mM MES, pH 6.5, 75 mM NaCl, 5 mM MgCl_2_, 0.02 % sodium azide and measured at 278 K.

For titrations 100 μM ^15^N-labelled TRIM25 407-630 or 439-630 were titrated with stock solutions of 10-20 mM natural abundance RNA or 250-300 μM RIG-I CARDs (aa 1-203 or 1-209) to a molar excess of 2.5-3 (details indicated in the figures). At each titration point a ^1^H-^15^N-HSQC spectrum was acquired. Spectra were processed using NMRPipe (36), visualized and peak shifts tracked using SPARKY (37). Peak shifts for PRY/SPRY 439-630 were assigned based on a previously published assignment (34). For PRY/SPRY 407-630, peak assignments were transferred from PRY/SPRY 439-630 and confirmed and extended, using HNCA and HNCACB backbone assignment experiments using CCPNMR (38). Secondary structure and order parameters were estimated from chemical shifts using TALOS and Sparta+ (39,40). R_1_ and R_2_ relaxation rates and ^1^H/^15^N heteronuclear NOEs were recorded using standard pulse sequences and analysed using PINT (41–43). Relaxation delays of 20, 50, 100, 150, 250, 300, 400, 500, 650, 800, 1000, 1300, 1600, 2400 ms for the R_1_ experiment and 16, 32, 48, 64, 80, 96, 128 ms for the R_2_ experiment were used.

### Isothermal titration calorimetry

Isothermal titration calorimetry (ITC) data was collected on a Malvern MicroCal PEAQ-ITC at 20°C in 20 mM MES, pH 6.5, 75 mM NaCl and 0.5 mM TCEP. Depending on the protein construct, RNA was titrated from the syringe at concentrations ranging from 20-860 μM into the cell containing protein with concentrations between 2-150 μM while stirring at 750 rpm (see Figure 2 and Supplementary Figure S3). For experiments with DENV-SL the RNA was kept in the cell at 15 μM and the protein titrated from the syringe at 110 μM. Experiments were typically done in triplicates and analysed using the MicroCal PEAQ-ITC analysis software. Data was fitted by assuming the simplest binding mode necessary to fully explain the data (one or two binding sites).

**Figure 2.**
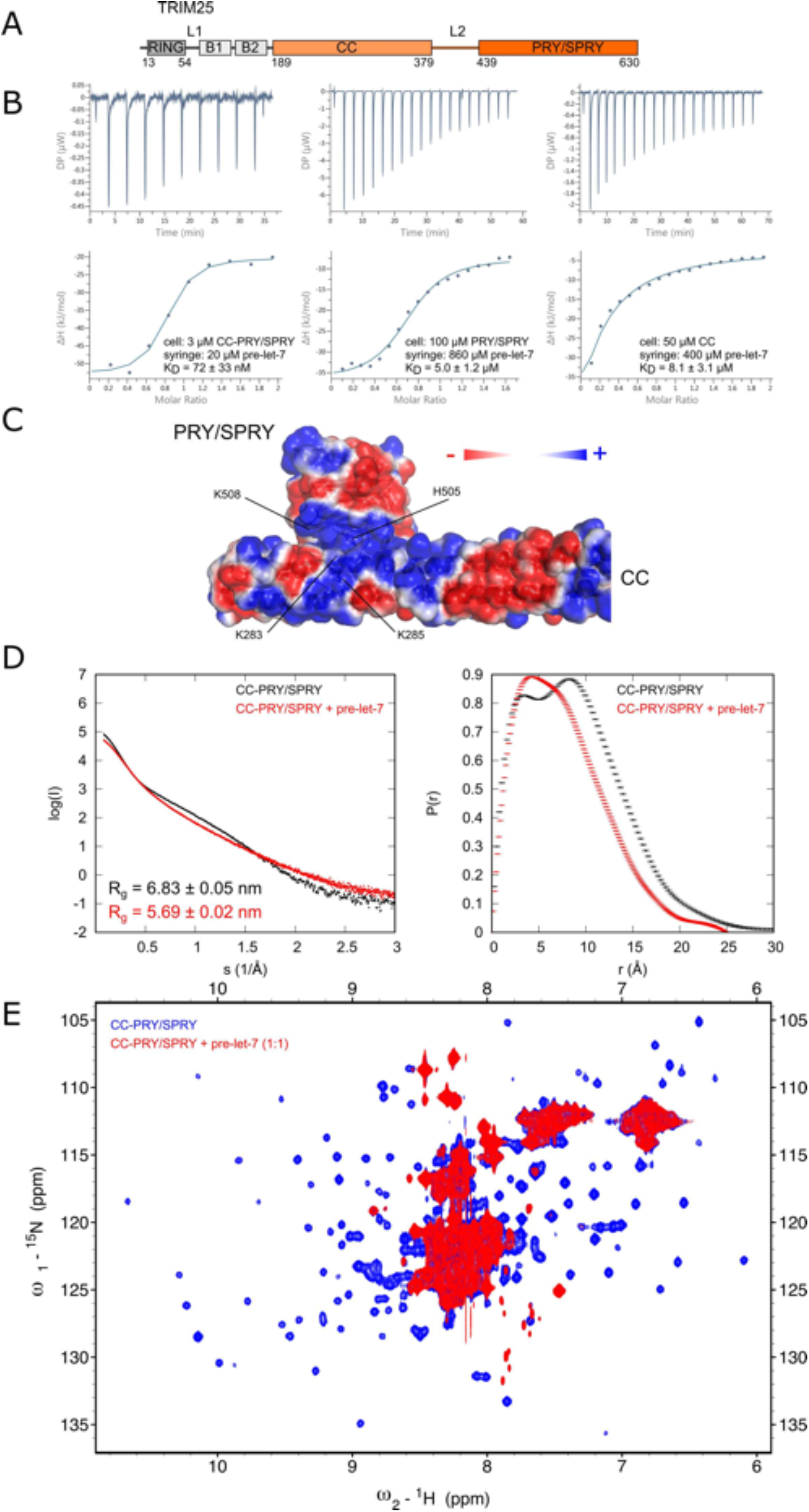
CC and PRY/SPRY domains bind RNA cooperatively: (**A**) Experiments in this panel were done with constructs of the CC and PRY/SPRY either in isolation or connected by the disordered L2 linker. **(B)** Binding isotherms of TRIM25 CC-PRY/SPRY, and PRY/SPRY titrated by pre-let-7 show that the PRY/SPRY alone is not sufficient to explain the nanomolar affinity of the longer CC-PRY/SPRY construct. The CC binds pre-let-7 similarly to the PRY/SPRY with micromolar affinity. The thermodynamic parameters of all ITC measurements can be found in Table 1. **(C)** The surface potential of the CC dimer with bound PRY/SPRY (PDB:6FLN) shows that they share a positively charged interface in close proximity to RNA binding site 1 marked by H505 and K508. **(D)** SAXS curves and pairwise distance distributions for the free TRIM25 CC-PRY/SPRY (black) and its complex with pre-let-7 RNA (red). The distance distribution of the free protein features two maxima, indicating independence of the two domains, whereas the distribution for the RNA-bound complex is much narrower and contains only one peak, indicating a conformational change towards a more compact form. **(E)** ^1^H,^15^N-HSQC spectra comparing the CC-PRY/SPRY in absence and presence of pre-let-7. The severe signal loss of peaks corresponding to residues in structured regions of the PRY/SPRY domain upon addition of equimolar amounts of pre-let-7 indicates that RNA keeps the PRY/SPRY domain at the CC interface leading to joint tumbling and thus increased transverse relaxation.

### Small-angle X-ray scattering

SAXS data were collected at the beamline P12, operated by EMBL Hamburg at the PETRA III storage ring (DESY, Hamburg, Germany) (44). Measurements were done at 20°C in 20 mM MES, pH 6.5, 75 mM NaCl and 0.5 mM TCEP in flow cell mode and a wavelength of 1.24 Å. For each sample and buffer 20-100 frames with 0.05-0.195 s exposure time were acquired and frames showing radiation damage manually removed. SEC-SAXS data was collected at the BM29 BioSAXS beamline, ESRF Grenoble using a wavelength of 0.99 Å, a Superdex 10/300 S200 Increase size exclusion column and analysed using Chromixs (45,46). Per run 2500 frames with 1s exposure time were collected. Data analysis was done using the ATSAS package version 2.8.3 (47). PRIMUS was used for frame averaging and buffer subtraction (48). The radius of gyration, *R_g_*, was estimated using the Guinier approximation in PRIMUS. Pair-wise distribution functions were calculated using GNOM (49). Collection statistics for SAXS measurements are summarized in Supplementary Table S1.

### Circular dichroism spectroscopy

CD spectra were acquired using a Jasco 815 circular dichroism spectrometer. Samples were measured at 20°C in quartz cuvettes with 1 mm path length in 20 mM MES, pH 6.5, 75 mM NaCl and 0.5 mM TCEP at 0.25 mg/ml.

### Filter binding assays

Filter binding assays were carried out in 200 μl of binding buffer (20 mM MES, pH 6.5, 75 mM NaCl, 0.5 mM TCEP). Structured RNA probes were refolded prior to experiments by heating at 85°C for 3 min and slow cooling to room temperature. A concentration series of TRIM25 CC-PRY/SPRY was incubated with 5′^32^P-labelled RNA for 10 min on ice and the samples were filtered through Whatman 0.45μm nitrocellulose filters. The protein/RNA complex was retained on the filters and detected by scintillation counting. Binding curves were fitted using SciDavis as:

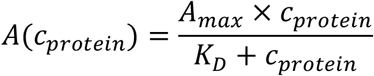

Where A_max_ and A(*c_protein_*) are measured activities. This approach only gives valid results, if the K_D_ is much smaller than the concentration of radio-labelled RNA. In cases, where this assumption was false the K_D_ was corrected as follows:

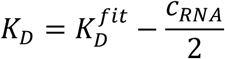

### Polynucleotide kinase (PNK) assays

HEK293T cells were cultured in DMEM with 10% FBS and antibiotics and seeded in 10cm plates (3×10^6^ cells per well) 24 hours prior transfection. Cells were transfected using Fugene transfection reagent and 10μg of HA-TRIM25 full-length WT or mutant in pcDNA3. Cells were washed with PBS, UV-crosslinked at 150 mJ/cm^2^ and lysed by sonication in lysis buffer (50mM Tris-HCl pH7.5, 100mM NaCl, 0.1% SDS, 1 mM MgCl_2_, 0.1 mM CaCl_2_, 1% NP40, 0.5% sodium deoxycholate, EDTA-free protease inhibitor). The cleared lysates were treated with 8 ng/ml of RNase A and 2U/ml Turbo DNase for 5 min at 37° C and immunoprecipitated for two hours using anti-HA magnetic beads (901502, BioLegend). The beads were washed three times with lysis buffer and three times with PNK buffer (50mM Tris-HCl, pH7.5, 50mM NaCl, 10mM MgCl_2_, 0.5% NP-40) at room temperature for three minutes respectively. The washed beads were incubated with 0.1mCi/ml [γ-^32^P] rATP, 1 U/ml T4 PNK in 5x NEB PNK buffer for 15 min at 37° C. After washing the samples with PNK buffer, the protein was eluted from the beads in low pH (0.1 M glycine, pH 2.0), neutralized with 0.2M Tris-HCl pH8.5 and separated on SDS-PAGE. The protein content of the gel was transferred to a nitrocellulose membrane, autoradiography recorded overnight using a phospho-imaging system and protein bands detected by Western blot using anti-TRIM25 (ab167154, Abcam, 1:2000).

### In cell ubiquitination assays

In cell ubiquitination of RIG-I CARDs by TRIM25 and its mutants was assessed as previously described (34). In short, HEK293T cells were cultured in DMEM with 10% FBS and antibiotics and seeded in 6-well plates (5×10^5^ cells per well) 24 hours prior to transfection. Cells were transfected using Fugene transfection reagent and 2μg DNA of the respective expression constructs in a pcDNA3 expression vector (1500 ng of RIG-I 2-200 with N-terminal FLAG-tag and 500 ng of TRIM25 full-length WT or mutants) For control samples equivalent amounts of pcDNA3 without insert were used instead of TRIM25 or RIG-I expressing plasmids.

24h after transfection cells were lysed in IP-lysis buffer (50mM Tris, 150mM NaCl, 5mM MgCl_2_, 1mM EDTA, 0.5% IGEPAL, protease inhibitors, 50nM NEM). For proteasome inhibition, cells were treated with 500nM carfilzomib 24 hours after transfection and lysed as before after additional 20 hours. An aliquot of the lysates was used for western blot to test and quantify the overexpression of TRIM25 (ab167154, Abcam, 1:2000) and FLAG-RIG-I CARDs (ab F3165, Sigma-Aldrich, 1:10.000) normalized to α-tubulin as loading control (T6199, Sigma Aldrich, 1:1000).

Immunoprecipitation (IP) using anti-FLAG magnetic beads (ab F3165, Sigma-Aldrich) was performed using similar amounts of lysate as quantified by BCA assays. IP was done to enrich overexpressed FLAG-RIG-I CARDs to investigate their ubiquitination state. IP eluates were used for western blot against FLAG-tag and ubiquitin (P4D1, Santa-Cruz Biotechnology, 1:500). Experiments were done at least in triplicates. Ratios of ubiquitinated to unmodified CARDs were quantified, normalized to their counterpart in the lane with no exogenous TRIM25 expression, averaged and the standard deviation calculated.

### Structure modelling

Models of the TRIM25 CC-PRY/SPRY complex with pre-let-7 were generated in CNS (50). As protein starting structures several previously published models that agree well with the solution structure of CC-PRY/SPRY were used (34). A model of pre-let-7 was generated using the SimRNAweb server (51). The two RNA molecules were randomly rotated and moved 100 Å away from the proteins and then pulled to the proteins using a rigid body minimization with ambiguous distance restraints to either the residues mapped by NMR or to any arbitrary protein residue. The resulting structures were subject to a short energy minimization by a Cartesian dynamics simulated annealing protocol (52). Pools of 200 structures each were calculated and fitted against the experimental SAXS data using CRYSOL (47).

## RESULTS

### Two distinct sites of the TRIM25 PRY/SPRY domain bind to single- and double-stranded RNA

Since RNA-binding of TRIM25 is structurally uncharacterized, we performed NMR titrations with the TRIM25 PRY/SPRY domain (Figure 1A) and the reported RNA target pre-let-7a-1 (Figure 1B). A minimal pre-let-7a-1@2 construct (pre-let-7 in the following, Supplementary Figure 1A) has been described to promote TRIM25-mediated regulation of let-7 by Lin28a and TUT4 (29). Upon addition of pre-let-7, we observed strong chemical shift perturbations (CSPs) clustered around two regions of the PRY/SPRY domain (Figure 1B-D). The first of the two binding sites (aa 456-511, binding site 1 in the following) is located at the C-terminus of the PRY motif with the strongest affected residues H505 and K508 in the flexible loop connecting ß-strands 3 and 4 (Supplementary Figure S1A). This site is located in close proximity to the interaction site between the PRY/SPRY and CC reported in Koliopoulos *et al*. (34). Our data thereby confirm the RNA-binding region (aa 470-508) observed using RBDmap by Castello *et al*. (7), but increase the accuracy to residue resolution. The second binding site (aa 549-605, binding site 2 in the following) with the strongest affected residues Y601 and K602 is located in a region formed by ß-strands 10 and 11, close to the N-terminal helix α1 (Supplementary Figure S1A). Although this region was not shown earlier to be involved in RNA binding, a central role in the recruitment of RIG-I has been reported (53). At this stage, we could not assess, whether the CSPs in this region are due to direct interaction with RNA or due to an allosteric effect.

Pre-let-7 is predicted to form a stem-loop structure (see Supplementary Figure S1B for a summary of predicted structures of all RNAs used in this study). The formation of double-stranded regions indicating stem formation has been validated by peaks in the imino region of ^1^H/^1^H-NOESY experiments (Supplementary Figure S1C). To further assess RNA specificity of these binding sites we designed shorter RNA constructs consisting of only the loop or stem of pre-let-7 (Supplementary Figure S1B). Upon titration with the single-stranded part of pre-let-7 (pre-let-7 loop), we observed CSPs only for residues located at binding site 1, whereas binding site 2 was unaffected (Figure 1E). To ensure that the short stem is double-stranded, we fused the strands together by a three bases long linker (pre-let-7 stem) and confirmed its stem formation in solution by 2D ^1^H,^1^H NOESY NMR (Supplementary Figure S1C). Titration with the resulting RNA construct or a 28-mer duplex RNA previously reported to bind TRIM25 (32) strongly affected only binding site 2 (Figure 1F, S1D). Together, this suggests different specificities for the two binding sites with binding site 1 binding single-stranded RNA and binding site 2 double-stranded RNA.

### TRIM25 binds RNA with high affinity and structure- and sequence-specificity

Previous research failed to uncover a clear RNA binding motif for TRIM25 (31). The presence of two binding sites with distinct binding preferences in close proximity suggests a possible binding selectivity for structural elements rather than sequence specificity. Indeed, comparing other reported RNA targets of TRIM25, we found that all feature stem loops of similar size and sequence as pre-let-7 (4,33). We used ITC to measure the affinity of several of these stem loops to TRIM25 CC-PRY/SPRY (Table 1, Figure 2A). We observed very tight binding of this construct to pre-let-7 (K_D_ =72 ± 33nM, Figure 2B) and similar stem loops (Table 1). Interestingly, the highest affinity measured was for a stem loop derived from the Dengue virus subgenomic RNA (DENV-SL) (4) that binds TRIM25 CC-PRY/SPRY with a K_D_ of 15.2 ± 3.0 nM (Table 1, Supplementary Figure S2A, left panel). TRIM25 CC-PRY/SPRY also binds one of the stem loops of the reported target long-noncoding RNA (lncRNA) Lnczc3h7a (33) with a K_D_ of 486 ± 77 nM (Supplementary Figure S2A, right panel). For pre-let-7 and to a lesser extent for the other stem loops we observed a binding isotherm suggesting the presence of several binding sites and a complex binding mode (Supplementary Figure S2B). Due to these complications, we could only quantify and compare the highest affinity binding event. The significance of the lower affinity binding events is not clear, but since they generally feature low N-values, they may represent RNA-induced oligomerisation. This is supported by the observation of protein aggregation upon RNA addition for most of these RNAs. Filter-binding assays confirmed the stronger binding of DENV-SL compared to pre-let-7 (Table 2, Supplementary Figure S2C). Affinities measured by filter-binding assays were generally higher than those by ITC, possibly because filter binding is more sensitive to aggregation.

**Table 1.** ITC data for various TRIM25 constructs and RNAs: Titrations with CC-PRY/SPRY constructs required fitting of a two-site model for larger RNAs. CC-PRY/SPRY and pre-let-7 showed several binding sites, of which only the highest affinity binding could be measured. The individual domains CC and PRY/SPRY show low N values, most likely explained by binding of several domains to one RNA. In rows with two values, two K_D_ values have been fitted to the binding curve. All experiments were done in triplicates and averaged values are reported.

| Protein | RNA | $K_D$ (M) | N | $\Delta H$ (kJ/mol) |
| --- | --- | --- | --- | --- |
| PRY/SPRY | pre-let-7-1a@2 | $5.0 \pm 1.2 \cdot 10^{-6}$ | $0.711 \pm 0.020$ | $-29.9 \pm 1.6$ |
| | DENV-SL | $466 \pm 130 \cdot 10^{-6}$ | $0.259 \pm 0.006$ | $-10.2 \pm 0.4$ |
|  | Lnczc3h7a-SL | very weak binding |  |  |
| | pre-let-7 loop | $42 \pm 53 \cdot 10^{-6}$ | $0.79 \pm 0.20$ | $-4.6 \pm 5.3$ |
|  | Lnczc3h7a loop | No binding detected |  |  |
| CC | pre-let-7-1a@2 | $8.1 \pm 3.1 \cdot 10^{-6}$ | $0.287 \pm 0.036$ | $-52 \pm 12$ |
|  | DENV-SL | n.d. due to phase-separation |  |  |
| | Lnczc3h7a-SL | $14 \pm 16 \cdot 10^{-6}$ | $1.1 \pm 0.5$ | $-3 \pm 2$ |
| | pre-let-7-1a@2 loop | $30 \pm 2 \cdot 10^{-6}$ | $1.28 \pm 0.02$ | $-2.8 \pm 0.2$ |
| | Lnczc3h7a loop | $32 \pm 2 \cdot 10^{-6}$ | $1.2 \pm 0.02$ | $-5.7 \pm 0.3$ |
| CC-PRY/SPRY | pre-let-7-1a@2 | $72 \pm 33 \cdot 10^{-9}$ | $0.750 \pm 0.028$ | $-33.0 \pm 2.3$ |
|  | pre-let-7-1a@2 stem-only | very weak binding |  |  |
| | pre-let-7-1a@2 loop-only | $4.1 \pm 1.9 \cdot 10^{-6}$ | $0.809 \pm 0.062$ | $-48.9 \pm 7.6$ |
| | DENV-SL | $15.2 \pm 3.0 \cdot 10^{-9}$<br>$106 \pm 35 \cdot 10^{-9}$ | $0.780 \pm 0.020$<br>$0.080 \pm 0.020$ | $-32.4 \pm 3.4$<br>$-330 \pm 100$ |
| | Lnczc3h7a-SL | $486 \pm 77 \cdot 10^{-9}$<br>$8.9 \pm 4.5 \cdot 10^{-6}$ | $1.25 \pm 0.05$<br>$0.104 \pm 0.063$ | $38.8 \pm 5.9$<br>$-17 \pm 82$ |
| CC-PRY/SPRY 381-392 7KA | pre-let-7-1a@2 | $428 \pm 55 \cdot 10^{-9}$<br>$967 \pm 11 \cdot 10^{-6}$ | $0.765 \pm 0.028$<br>$0.13 \pm 0.22$ | $-17.0 \pm 1.2$<br>$-270 \pm 410$ |
| CC-PRY/SPRY H505E K508E | pre-let-7-1a@2 | $484 \pm 48 \cdot 10^{-9}$<br>$4.19 \pm 0.50 \cdot 10^{-6}$ | $0.972 \pm 0.030$<br>$0.196 \pm 0.043$ | $-10.2 \pm 0.76$<br>$-91 \pm 18$ |
| CC-PRY/SPRY K602E | pre-let-7-1a@2 | $196 \pm 22 \cdot 10^{-9}$<br>$2.54 \pm 0.12 \cdot 10^{-6}$ | $1.08 \pm 0.03$<br>$0.050 \pm 0.043$ | $-3.31 \pm 0.53$<br>$-270 \pm 230$ |
| CC-PRY/SPRY K283A K285A | pre-let-7-1a@2 | $606 \pm 124 \cdot 10^{-9}$<br>$16.6 \pm 2.1 \cdot 10^{-6}$ | $1.36 \pm 0.06$<br>$0.12 \pm 0.20$ | $-5.53 \pm 0.90$<br>$-145 \pm 215$ |
| CC-PRY/SPRY<br>H505E K508E<br>K602E | pre-let-7-1a@2 | $790 \pm 160 \cdot 10^{-9}$<br>$132 \pm 24 \cdot 10^{-6}$ | $0.973 \pm 0.025$<br>$0.06 \pm 0.29$ | $-2.83 \pm 0.49$<br>$-300 \pm 1400$ |
| CC-PRY/SPRY<br>K283A K285A<br>H505E K508E<br>K602E | pre-let-7-1a@2 | $1.32 \pm 0.37 \cdot 10^{-6}$<br>$124 \pm 28 \cdot 10^{-6}$ | $0.972 \pm 0.030$<br>$0.26 \pm 0.12$ | $-0.2 \pm 1.7$<br>$-210 \pm 130$ |
| | DENV-SL | $363 \pm 84 \cdot 10^{-9}$ | $0.466$<br>$\pm 0.008$ | $14.17 \pm 0.46$ |
|  | Lnczc3h7a-SL | No binding detected |  |  |
| CC-PRY/SPRY<br>Y463S Y476S | pre-let-7-1a@2 | $252 \pm 61 \cdot 10^{-9}$<br>$17 \pm 11 \cdot 10^{-6}$ | $1.42 \pm 0.03$<br>$0.05 \pm 0.10$ | $-8.6 \pm 1.2$<br>$-290 \pm 580$ |

**Table 2.** Filter-binding assays confirm binding to stem-loops

| protein | RNA | K<sub>D</sub> (M) |
| --- | --- | --- |
| CC-PRY/SPRY | pre-let-7-1@2a | $10.2 \pm 3.8 \cdot 10^{-9}$ |
| | DENV-SL | $1.63 \pm 0.38 \cdot 10^{-9}$ |
| | RoX2 SL6 | $18.1 \pm 2.2 \cdot 10^{-9}$ |

Because of this strong interaction with several stem-loops of similar size, but limited sequence similarity, we hypothesized that TRIM25 specifically recognizes stem-loop structures. To confirm this hypothesis, we tested binding to a stem loop that is similar in size but has little sequence similarity and is functionally unrelated. We chose stem-loop 7 of *Drosophila* lncRNA roX2, due to its origin in an evolutionary distantly related organism and its involvement in dosage compensation, an unrelated function (54). Filter-binding assays found an interaction with nanomolar affinity supporting the hypothesis that specificity of RNA binding of TRIM25 is largely mediated by RNA structure rather than sequence (Table 2, Supplementary Figure S2B). However, this does not rule out that individual binding sites have sequence-specific motifs that further contribute to selective RNA binding. However, this possibly remained undetected by previous approaches due to failure to deconvolute the binding contribution of distinct binding sites to specificity and take into account structural requirements of the RNA. As an example, we found that the lnczc3h7a loop showed only very weak CSPs on binding site 1 of the PRY/SPRY when compared to pre-let-7 loop (Supplementary Figure S1). By ITC the affinity for lnczc3h7a loop to the PRY/SPRY was too weak to be quantified (Supplementary Figure S2D, right panel) while for pre-let-7 loop robust, but weak binding was found K_D_ = 42 ± 53 μM, Table 1, Supplementary Figure S2D, left panel). This suggests that sequence recognition by the single-stranded binding sites on the PRY/SPRY additionally contributes to binding specificity and explains the observed reduction in affinity for the U-rich lnczc3h7a-stem-loop when compared with the A/G-rich pre-let-7 and DENV-SL-loops and is in line with previous findings that suggested a preference for G and A-enriched sequences (29).

### Cooperative binding of PRY/SPRY and CC domain to RNA

As reported previously, we found that the affinity of the PRY/SPRY domain for pre-let-7 RNA (5.0 ± 1.2 μM, as measured by ITC, Table 2, Figure 2B) is much weaker than that of CC-PRY/SPRY (31,32). The RNA-binding of the PRY/SPRY domain alone is therefore not sufficient to explain the high affinity binding of TRIM25. The complex binding isotherm suggests that an additional binding site might be necessary for high affinity binding (Supplementary Figure S2A). This is also supported by the observation that the strong increase of affinity in the presence of the CC domain and linker region is only observed for larger stem-loop RNAs, while the shorter isolated stem and loop constructs show binding isotherms that can be explained by a single binding site model and only slightly stronger binding to CC-PRY/SPRY than to single domains is observed (Table 1, Supplementary Figure 2E). Sanchez *et al*. (32) described a lysine-rich stretch in the L2 linker connecting the CC and PRY/SPRY domains as necessary for binding of double-stranded RNA. Mutation of these lysines to alanine (381-392 7KA) reduced the affinity to double-stranded RNA 20-fold in the original study (32). However, in our experiments this mutant had weaker effect on the binding of short stem loop RNAs to CC-PRY/SPRY K_D_ =428 ± 55 nM, Supplementary Figure S2F), making it unlikely that this binding site alone is responsible for the observed increase in affinity.

We noted that in the crystal structure of the CC-PRY/SPRY dimer (PDB:6FLN) binding site 1 is part of a larger positively charged surface that extends onto the CC (Figure 2C). Together, the PRY/SPRY domain and CC could form a joint interaction surface that would allow for positive cooperativity (synergistic binding) and could explain the tighter binding of RNA by the CC-PRY/SPRY construct as observed by ITC and filter binding assays. The CC is not suitable for NMR studies due to its size and extended conformation, which causes slow molecular tumbling and NMR resonance line broadening beyond detection. Instead, we used ITC to test the CC’s putative RNA-binding capacity (14,55) (Table 1). We found that the CC binds pre-let-7 with low micromolar affinity (8.1 ± 3.1μM) (Figure 2B), slightly weaker than the PRY/SPRY domain. The more than 500-fold increase in affinity of the CC-PRY/SPRY construct over the isolated domains is a hallmark of chelating cooperativity, in which multiple, individually weak binary interactions cooperatively lead to the formation of a stable multimeric complex (56,57). We have previously shown the existence of a transient interaction between the CC and PRY/SPRY domain in solution, that despite its low affinity is critical for the E3 ligase activity of TRIM25 (34). Chelating cooperativity also offers a possible explanation for the observed additional binding events for CC-PRY/SPRY in ITC, as these cooperative effects are concentration dependent and above a critical concentration the binding mode switches from positive cooperativity to non-cooperative binding, leading to interactions of the single binding sites with distinct RNA molecules (56). This RNA-mediated chelating cooperativity in the CC-PRY/SPRY construct suggests a mutual stabilisation of this otherwise transient domain-domain interaction. We therefore used solution small-angle X-ray scattering (SAXS) to test for a stabilisation of the transient interaction between CC and PRY/SPRY domains through RNA-binding. A comparison of the SAXS curves of free TRIM25 CC-PRY/SPRY and the complex with pre-let-7 shows a significant decrease in the radius of gyration upon RNA-binding (R_g_ = 6.83 ± 0.05 nm for the free protein compared to Rg = 5.69 ± 0.02 nm for the complex), indicating that the complex adopts a more compact conformation than the free protein in solution (Figure 2D), where the PRY/SPRY domain is mostly detached from the CC domain (34). Similar effects were observed for the complex with a lnczc3h7a stem-loop (R_g_ = 5.81 ± 0.18 nm), confirming that this is a general property of stem-loop binding (Supplementary Figure S3A). The effect also occurs in SEC-SAXS with an excess of pre-let-7 clearly showing that it is not an artefact caused by the scattering contribution of the free RNA (Supplementary Figure S3B).

We then calculated an ensemble of CC-PRY/SPRY-RNA complex structures to investigate if this observed change in the radius of gyration is due to a conformational change of the protein or binding of the RNA closer to the centre-of-mass of the complex. As starting structures, previously published models of TRIM25 CC-PRY/SPRY were used that fit the experimental SAXS curve well (34). Then, we docked a model of pre-let-7 either to the interfaces on the PRY/SPRY observed by NMR or anywhere on the surface of the protein (Supplementary Figure S3C). All models showed radii of gyration that where significantly bigger than the obtained models (> 6.08 nm) and none of them fitted well to the experimental SAXS curve for the complex. This suggests that RNA binding alone cannot explain the observed decrease in R_g_ and additional conformational changes of the protein must occur, that bring the PRY/SPRY closer to the centre-of-mass of the entire protein.

This is also evident from the pairwise distance distribution, P(r), obtained from the SAXS data, where the free protein shows a broad distribution with two distinct maxima, indicating that the two domains tumble independently. In contrast, the distribution for the complex is shifted to smaller distances and shows only a single maximum, indicating that the complex tumbles as a single, compact entity (Figure 2D, Supplementary figure S3A).

To further confirm that this conformational change is indeed due to a stabilization of the transient CC:PRY/SPRY interaction through RNA binding, we monitored changes of TRIM25 CC-PRY/SPRY upon addition of pre-let-7 using NMR. In the absence of RNA, the HSQC of CC-PRY/SPRY is dominated by sharp, high intensity peaks in the centre corresponding to the disordered L2 linker region. Further well dispersed peaks overlap well with the spectra of the PRY/SPRY domain but show weaker intensity than in the spectra of the isolated domain (Supplementary Figure 3D). This agrees with the previously described transient interaction between CC and PRY/SPRY, as only the free, but not the CC-bound PRY/SPRY domains are observed. There is no overlap with spectra of the isolated CC, indicating that the large, extended CC dimer is too big to be observed in NMR. Upon addition of an excess of pre-let-7 in contrast to the CSPs for the isolated PRY/SPRY in the CC-PRY/SPRY construct the peaks of the PRY/SPRY disappear completely, while the peaks of the L2 linker remain visible. This suggests that the RNA-bound PRY/SPRY is no longer observable in NMR. This agrees with a stabilization of the CC:PRY/SPRY interaction and a shift of the equilibrium towards the CC bound state (Figure 2E).

We therefore conclude that RNA binding enhances the interaction between the CC and PRY/SPRY domains and leads to a more compact conformation of the protein.

### Mutational analysis confirms RNA-binding sites on PRY/SPRY and CC

To better understand the relative contributions of the PRY/SPRY RNA binding sites identified by NMR, and the binding site on the CC, inferred from the crystal structure of CC-PRY/SPRY (Figure 2B), we created point mutants and tested their effect on RNA binding using ITC (Figure 3A, B, Table 1). Mutation of binding site 1 residues (H505E/K508E) in the context of the CC-PRY/SPRY construct led to a 6-fold reduction of affinity to pre-let-7 compared to the WT (K_D_ = 484 ± 48 nM). On binding site 2, Y601 and K602 show the strongest CSPs and a K602E mutant reduced binding by a factor of 2.5 (K_D_ = 196 ± 22 nM). Combination of these mutants (triple-mutant H505E/K508E/K602E) further reduces RNA-binding more than ten-fold compared to wildtype (K_D_ = 790 ± 160 nM). This suggests that both binding sites on the PRY/SPRY domain are critical for cooperative binding. The effect of these mutants is however not as strong as that of previously published, less conservative deletions or mutation of entire regions (31,32).

**Figure 3.**
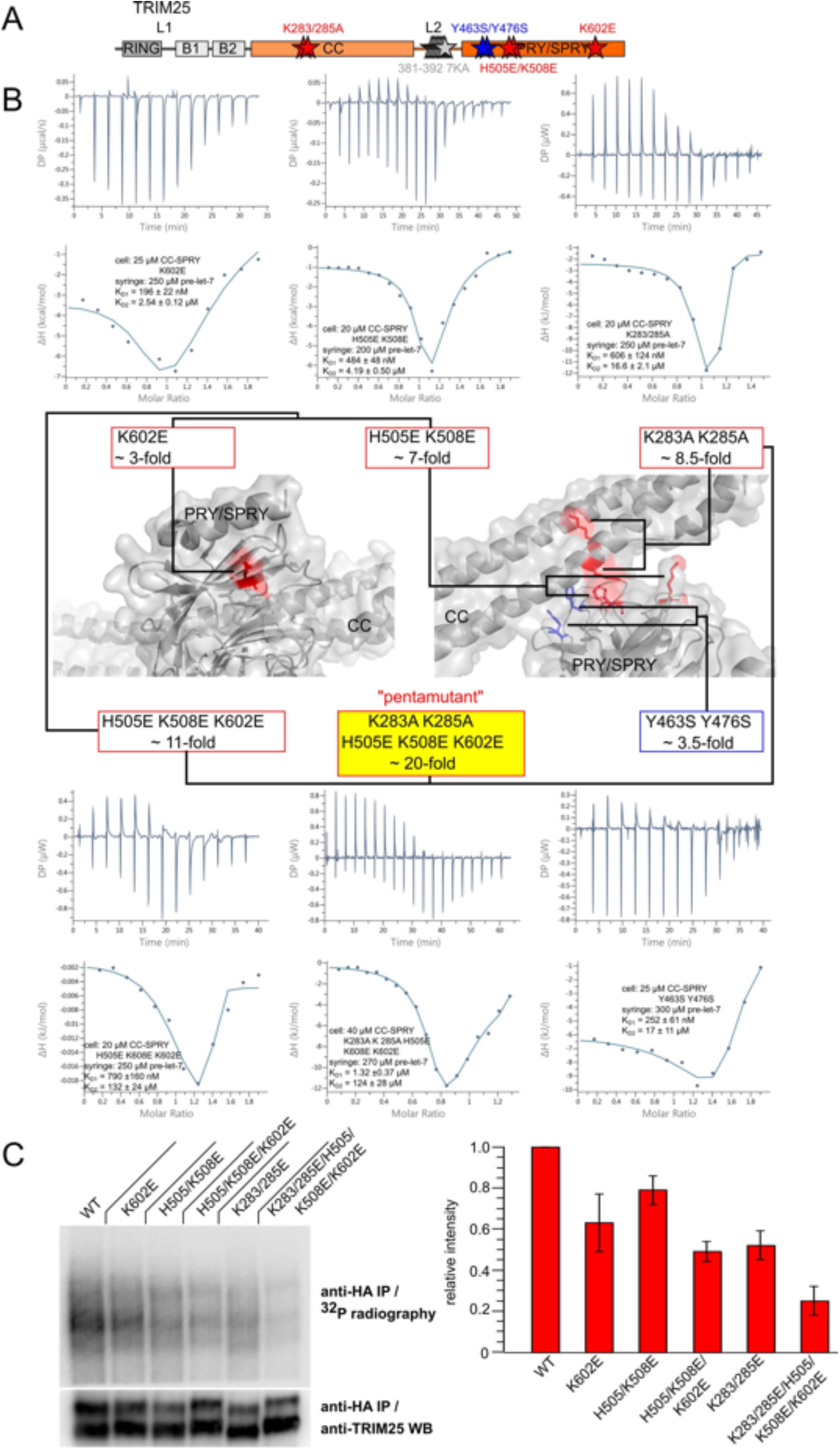
Mutational analysis of the RNA binding interface of TRIM25: **(A)** Point mutants affecting RNA binding of TRIM25 were found on both CC and PRY/SPRY. A reduction of RNA binding through the 381-392 7KA mutant in the disordered L2 linker was confirmed (Supplementary figure S2E)(32). **(B)** Isothermal titration calorimetry shows that mutants on both binding sites of the PRY/SPRY (H505E/K508E on binding site 1, K602E on binding site 2) and on the CC (K283A, K285A) reduces RNA binding in the context of the CC-PRY/SPRY construct. Crucially the mutants of binding site 1 on the PRY/SPRY and those on the CC form a joint interface allowing for cooperative binding of RNA. Combination of the mutants leads to a penta-mutant that reduces binding to pre-let-7 around 20-fold. In addition to these mutants of the RNA binding interface also the mutant Y463/476S previously described to reduce the CC:PRY/SPRY interaction affects RNA binding, further supporting the crucial role of this interaction for RNA binding (34). **(C)** RNA binding of TRIM25 is also confirmed in cells using polynucleotide kinase (PNK) assays. All mutants lead to a significant reduction of cellular RNA binding. The histogram shows quantification of cellular RNA binding from four replicates.

We were not able to map the full RNA binding interface on the CC due to technical limitations mentioned above and therefore relied on indirect information to design mutants of the CC based on the CC-PRY/SPRY structure. Thus, we inferred RNA binding residues from amino acids which are surface-exposed, close to the CC:PRY/SPRY interface and belong to typical RNA-binding residues like lysines, arginines and aromatic residues and found that a double mutant on the basic surface close to the PRY/SPRY binding site (K283A/K285A) in the context of the CC-PRY/SPRY reduced RNA-binding about 8-fold (K_D_ = 606 ± 124 nM, Figure 3B). It is however likely that additional residues on the CC are involved in RNA binding, since we restricted mutations to the proximity of the PRY/SPRY-binding site. Combination of the mutants on the CC and PRY/SPRY results in a penta-mutant that reduces binding to pre-let-7 almost 20-fold (K_d_ = 1.32 ± 0.37 μM). To confirm that the effect of these mutants is not limited to pre-let-7 we tested binding of the penta-mutant also to DENV-SL and the lnczc3h7a stemloop and observed a more than 20-fold reduction for the first (K_D_ = 363 ± 84 nM) and no longer detectable binding for the latter (Supplementary Figure S4A). We confirmed proper folding of the mutants to ensure that the observed effects are due to specific disruption of RNA binding interactions rather than misfolding of the domain: The ^1^H,^15^N-HSQC spectrum of the triple-mutant PRY/SPRY showed only negligible chemical shift perturbations compared to the wildtype, confirming proper folding of the domain (Supplementary Figure S4B). Of note, this was not the case for the previously reported RNA-binding deficient deletion mutant (31), for which the ^1^H,^15^N-HSQC spectrum only shows peaks in the centre, indicative of aggregation or an unfolded domain (Supplementary Figure S4C). Either way, the structural integrity of this deletion mutant is compromised, and RNA binding is likely impaired due to the absence of a folded PRY/SPRY domain. NMR spectra of the CC show only peaks corresponding to the flexible termini of this construct due to its large size and elongated architecture, which leads to fast transverse relaxation of structured regions (Supplementary Figure S4D). However, the CC domain strongly dominates the CD spectrum of the CC-PRY/SPRY construct due to its high α-helix content. Thus, we used CD spectroscopy to ensure that the K283/285A mutant on the CC does not impair folding in the context of the penta-mutant either (Supplementary figure 4E).

Interestingly, we found that the double mutant Y463S/Y476S, that reduces the interaction between CC and PRY/SPRY (34), also reduced RNA binding (K_D_ = 252 ± 61 nM, Figure 3B) although these residues were unaffected by RNA binding in our NMR titrations with the isolated PRY/SPRY domain. This further supports that the CC-PRY/SPRY interaction has a crucial role in RNA-binding and suggests a mutual stabilization of the domain/domain and domain/RNA interactions.

To validate RNA binding of TRIM25 in HEK293T cells we used polynucleotide kinase (PNK) assays. These confirmed that TRIM25 indeed binds RNA also in cells and that RNA binding mutants showed a significant reduction of binding to host RNAs (Figure 3C). Especially the penta-mutant led to a dramatic loss of RNA binding. This suggests that the RNA binding interface we describe here is not limited to a small number of model-RNAs but is also crucial for the interaction of TRIM25 with most or all of its host target RNAs. The set of point mutants therefore allows to assess the downstream consequences of loss of host RNA binding of TRIM25 without perturbations to the protein structure.

### RNA binding regulates ubiquitination activity

We have previously demonstrated the importance of the transient CC-PRY/SPRY interaction for the catalytic activity of TRIM25 in RIG-I signalling (34) and as we propose a stabilization of this interaction upon RNA binding, we next investigated the effect of RNA binding mutants on the ubiquitination activity of TRIM25. To do so, we co-transfected HEK293T cells with wildtype or mutant TRIM25 and FLAG-tagged RIG-I CARDs (Figure 4A, Supplementary Figure 5). Expression levels of TRIM25 and ubiquitination of the CARD domains were detected by Western blotting. Overexpression of wildtype TRIM25 caused robust ubiquitination of the RIG-I CARDs. In line with our hypothesis, both K602E and H505E/K508E on the PRY/SPRY as well as K283A/K285A on the CC and combinations thereof reduced ubiquitination of RIG-I CARDs, suggesting that reduced RNA binding affects E3 ligase activity in cells (Figure 4B). The penta-mutant shows a weaker phenotype than the individual mutants, despite being stronger impaired in RNA binding. This observation remains unexplained. It is noteworthy that the effect of the mutants is strongest for polyubiquitination, while mono-ubiquitination is less affected (Figure 4C). It is possible that mono-ubiquitination of CARDs is caused by an E3 ligase other than TRIM25 or RNA binding might be required for chain elongation, but not initiation. The latter could be an E2 specific effect, supported by the observation reported earlier that Ubc13/Uev1A, the E2 complex specific for the production of K63-linked ubiquitin chains *in vitro* only produces unanchored ubiquitin chains, suggesting that a second E2 is necessary for chain initiation (58,59).

**Figure 4.**
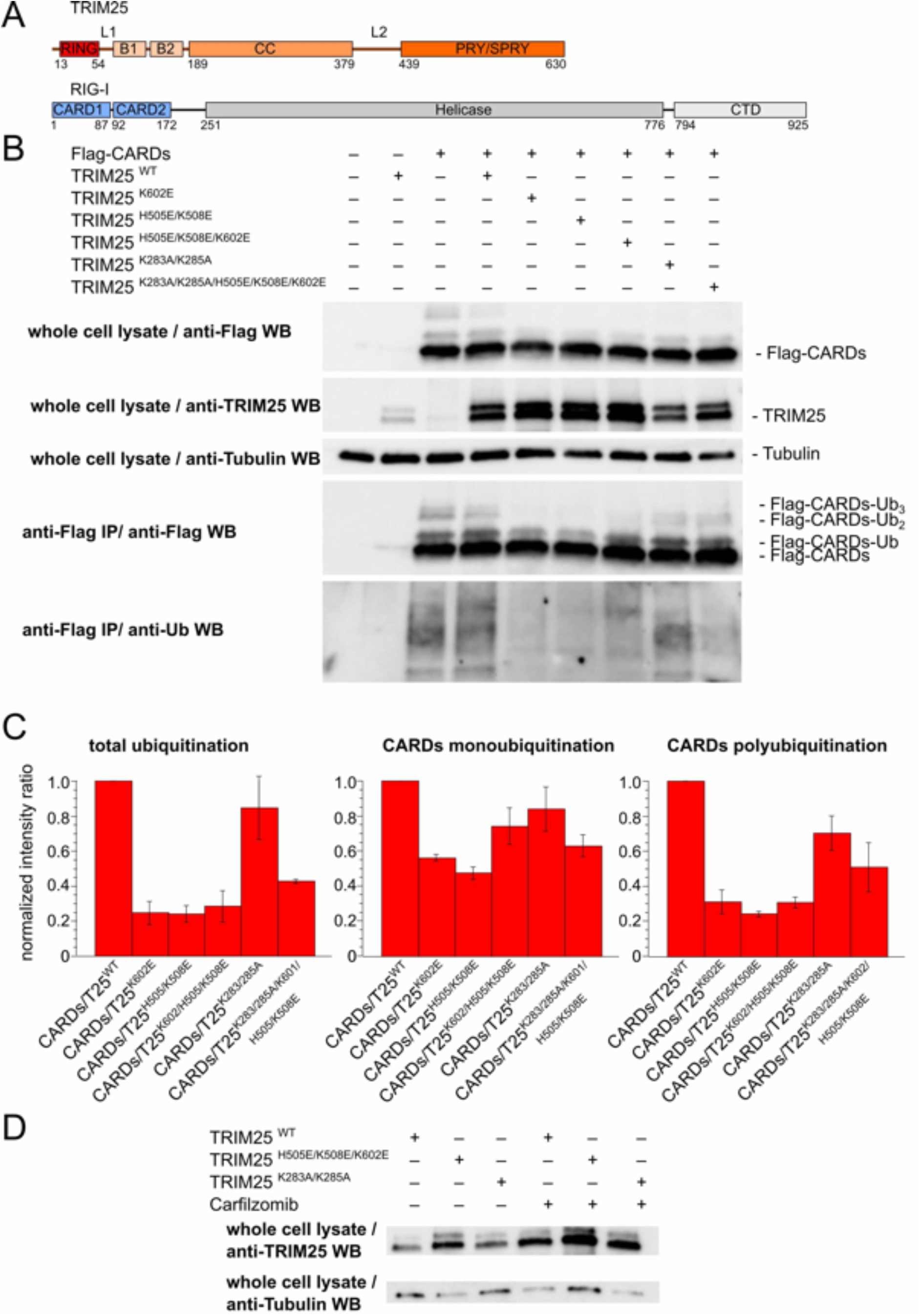
The RNA binding interface is required for RIG-I ubiquitination in cells: Isolated RIG-I CARDs with Flag-tag were overexpressed with TRIM25 WT or mutants **(A)**, immunoprecipitated and probed for ubiquitination **(B). (C)** Mutants of the RNA-binding interface reduce ubiquitination of RIG-I CARDs. Total ubiquitination was measured on anti-ubiquitin Western blots after IP against Flag, while mono- and polyubiquitination of RIG-I was measured on anti-Flag Western blots. Monoubiquitination appears to be less affected than polyubiquitination. **(D)** The mutants on the PRY/SPRY show increased expression levels compared to the wildtype in panel (A). Inhibition of the proteasome by carfilzomib stabilizes expression levels of both the WT and mutants, but the stabilization is weaker for mutants on the PRY/SPRY, suggesting that the difference is at least in parts due to reduced auto-ubiquitination.

The effect of the mutants is even more striking, given that, despite transfection of the same plasmid amount, mutants H505E/K508E and K602E on the PRY/SPRY, but not K283/285A on the CC, showed consistently elevated expression levels compared to the wildtype. RT-PCR showed no significant differences in mRNA levels between WT and mutants, suggesting that this difference is likely due to differences in protein stability. TRIM25 promotes proteasomal degradation of several of its substrates, such as MAVS, ZAP or 14-3-3σ through the production of K48-linked ubiquitin chains and is also known to RNA-dependently auto-ubiquitinate *in vitro* (18,31,58,60,61). To test if the difference in expression levels can be explained by differences in auto-ubiquitination and proteasomal degradation, we inhibited the proteasome using Carfilzomib. This stabilized both wildtype and H505E/K508E/K602E triple mutant TRIM25 but had a much more dramatic effect on the wildtype, leading to similar expression levels than the mutant (Figure 4D). This points towards a crucial role of TRIM25-RNA binding not only in RIG-I ubiquitination but also in auto-ubiquitination of TRIM25. However, as the RNA binding deficient mutants include the replacement of lysines, it cannot be excluded that the auto-ubiquitination target lysine of TRIM25 is removed. Nevertheless, together with the strong reduction in substrate ubiquitination, this suggests that RNA binding enhances auto-ubiquitination of TRIM25 in cells and could thereby regulate its proteasomal degradation and protein levels.

### RIG-I CARDs do not interact directly with TRIM25 PRY/SPRY domains

To gain further insight into the structurally so far uncharacterized interaction of TRIM25 and RIG-I, we attempted to use NMR to map the interaction site and determine the affinity. Previous studies have established that the RIG-I CARDs alone are sufficient for robust TRIM25 interaction and ubiquitination in cells and of TRIM25 only the PRY/SPRY is required for RIG-I binding (17,53) (Figure 5A). We therefore titrated unlabelled RIG-I CARDs into ^15^N-labelled TRIM25 PRY/SPRY to monitor expected chemical shift perturbations and/or line broadening to identify interacting residues. However, while such an approach would almost certainly detect any direct interaction with affinities down to low millimolar affinities, we did not observe any signs of interaction (Figure 5B). While not completely ruling out the possibility of a TRIM25-RIG-I interaction, this strongly suggests the absence of a direct interaction between TRIM25 PRY/SPRY and RIG-I CARDs. D’Cruz *et al.* (53) proposed a mechanism of RIG-I binding, that would involve displacement of the N-terminal helix α1 of the PRY/SPRY by a similar helix in the CARDs. We therefore characterized secondary structure and flexibility of α1 using ^15^N relaxation measurements and heteronuclear NOEs (HetNOE) (Figure 5C-E). We found that α1 forms a stable helix in solution and tumbles together with the core domain, as residues in this region have the same apparent rotational correlation time as the entire domain. This indicates that a mechanism that involves unfolding or flipping out of α1 is very unlikely and agrees with earlier observations, that although the functional interaction between TRIM25 and RIG-I is very well established in cells, there is no evidence for their direct interaction *in vitro* (32). Several additional factors thought to stabilize the interaction, including both proteins and RNAs, have been proposed and might resolve this discrepancy (33,55,62).

**Figure 5.**
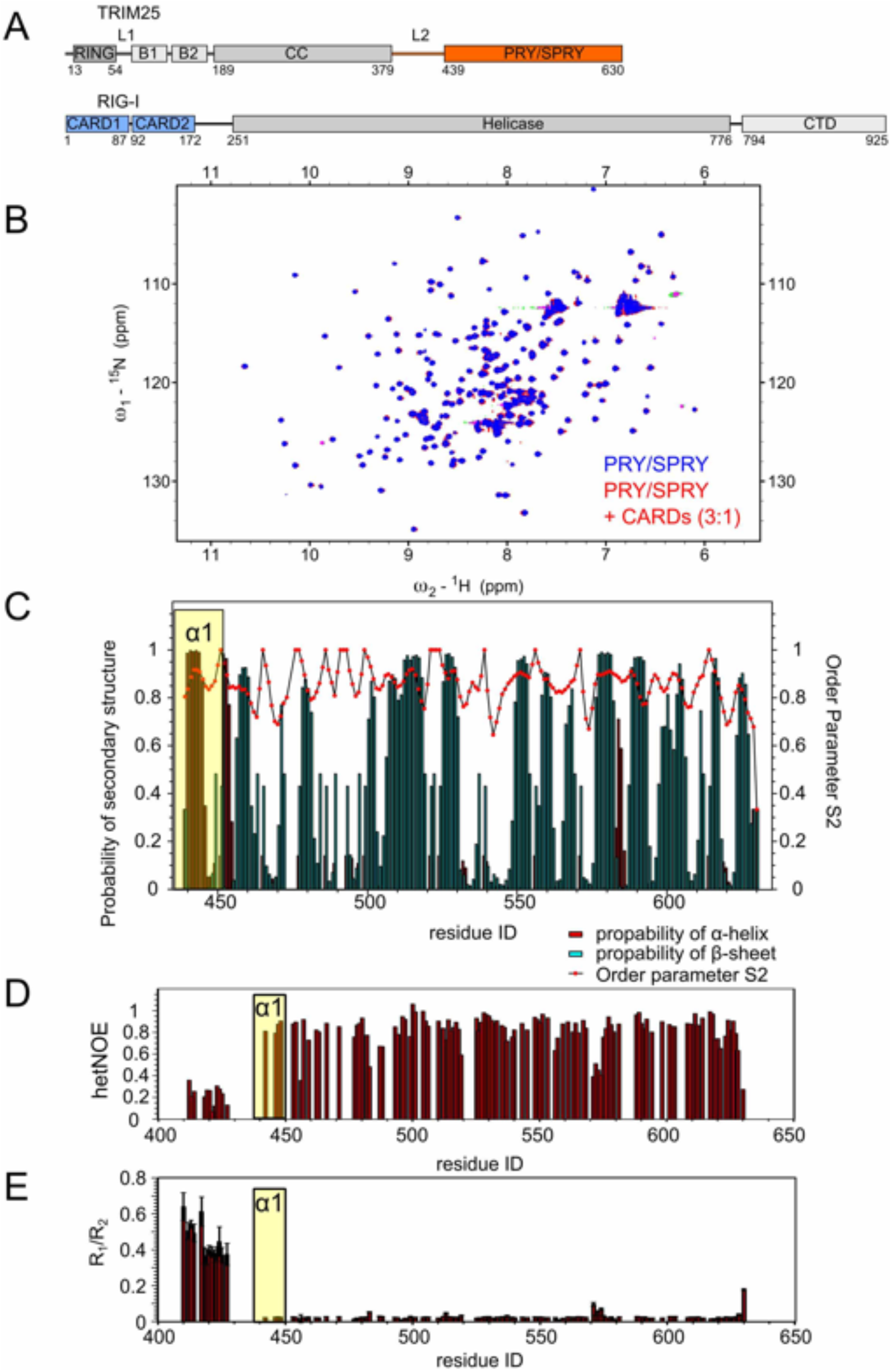
RIG-I interaction and dynamics of helix α1: **(A)** In cell the TRIM25 PRY/SPRY and RIG-I CARDs are sufficient to mediate the interaction between these proteins (17). We therefore focused on these domains for our investigation. **(B)** NMR titration of TRIM25 PRY/SPRY by RIG-I CARDs. No CSPs or changes in signal intensity, that would indicate a direct interaction were observed. **(C)** Secondary structure prediction and order parameter for the TRIM25 PRY/SPRY (aa 439-630) based on the previously published assignment (34). In agreement with the crystal structure (PDB:6FLM) the α1 region (marked in yellow) is predicted to form a helix in solution. This is supported by heteronuclear NOEs **(D)**, that show strong correlation of the backbone N-H vectors, confirming that α1 is formed. Relaxation measurements **(E)** give information on the backbone dynamics of individual residues. The residues in the α1 region show similar dynamics as the core domain. By contrast the residues belonging to the L2 linker (aa 407-435) are much more dynamic. D’Cruz et al. (53) proposed a mechanism of CARD binding, that would involve rearrangement of helix α1 by a similar helix in the CARD domains. This mechanism is not supported by our data.

## DISCUSSION

In summary, we show that TRIM25 achieves very tight binding of RNA through several binding sites on the CC and PRY/SPRY domain, with each binding site showing only weak affinities to RNA and different specificities. While we confirm and refine a previously identified binding region on the PRY/SPRY, we found additional binding sites on the PRY/SPRY and CC. The novel second binding site on the PRY/SPRY seems to be specific for double-stranded RNA and overlaps with a region previously thought to be involved in RIG-I binding (53). The close proximity of binding sites specific for single and doublestranded RNA suggests a specificity for structured RNAs and indeed we found binding with nanomolar affinity to several stem-loops. This may explain the failure of previous studies to identify a clear RNA motif for TRIM25 (29). On the other hand, by comparing the loop sequences of pre-let-7 and lnczc3h7a, we could also observe a preference for A/G-rich single-stranded motifs for binding site 1. The highly cooperative binding mode involving multiple binding sites on different domains is critical for the E3-ubiquitin ligase activity of TRIM25, as RNA binding stabilizes the weak interaction between the CC and PRY/SPRY domain which was previously shown to be important for TRIM25’s catalytic activity (34). In the absence of structural data of the complete tripartite motif of TRIM25 the importance of this interaction can be understood by considering two recently published structures of the tripartite motif of TRIM28, a distant relative of TRIM25 (63,64) (Figure 6A). These structures show that the RING domain binds the CC at a position close to the binding site of the PRY/SPRY domain in TRIM25 (Figure 6A). RNA-mediated binding of the PRY/SPRY domain to the CC will therefore bring the PRY/SPRY domain and any substrate bound by it in close proximity to the RING domain and the bound and ubiquitin-loaded E2 conjugating enzyme, facilitating substrate ubiquitination (Figure 6B). This is in accordance with the finding of us and others, that TRIM25 RNA-binding is required for efficient ubiquitination of the RIG-I CARDs (32). It is however not clear how RING dimerization, that is necessary for the E3 ligase activity of TRIM25 but not TRIM28, is achieved in this model (58,59,65). Such RING dimerization could occur between two TRIM25 dimers stacking up end to end. Alternative models have been proposed, that place the RING domain closer to the centre of the CC, allowing for RING dimerization within the TRIM25 dimer (59,66)

**Figure 6.**
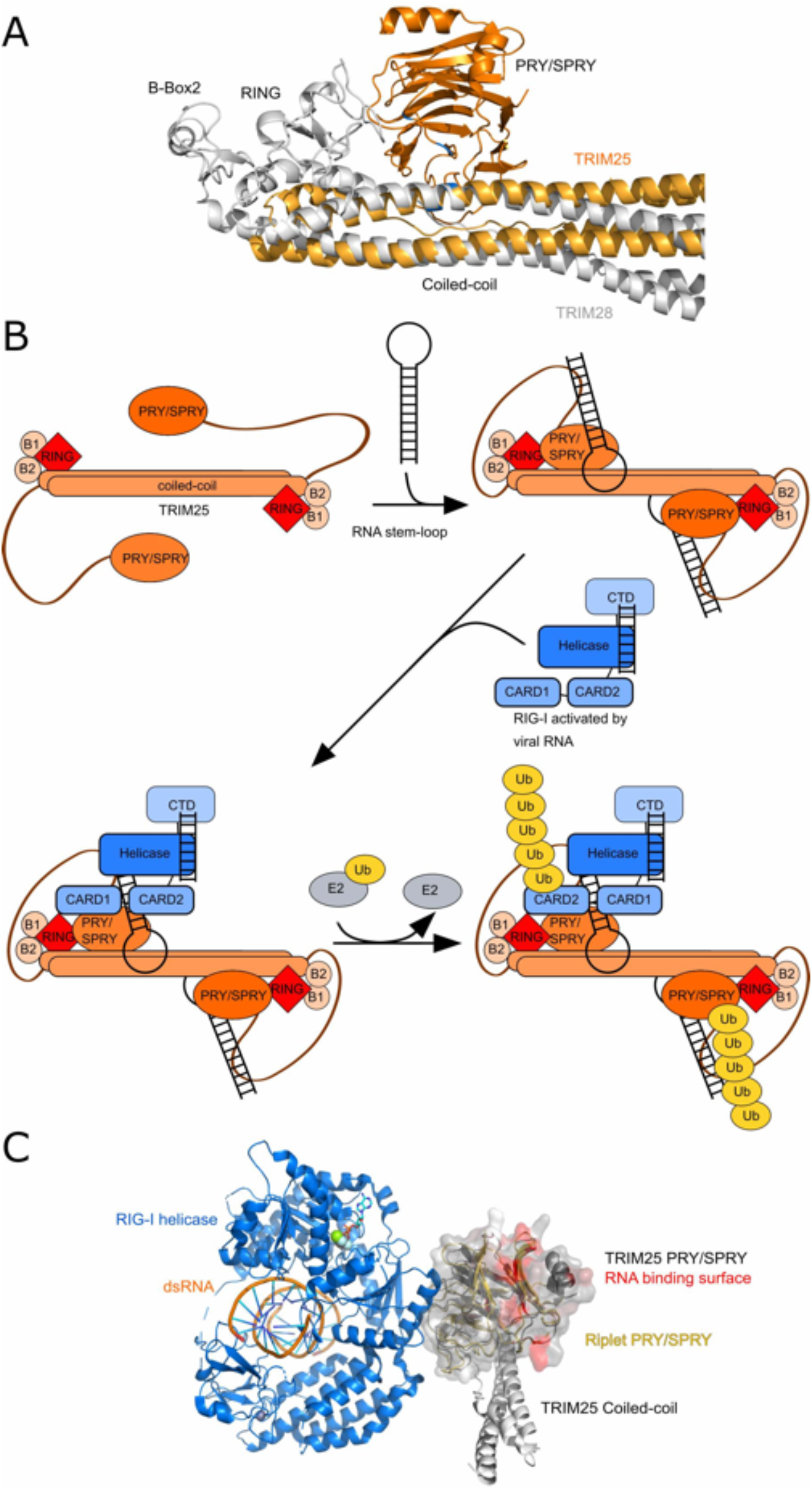
Potential model of full-length TRIM25 and RIG-I ubiquitination: **(A)** Alignment of the crystal structure of the TRIM28 tripartite motif (PDB: 6QAJ) with the TRIM25 CC-PRY/SPRY structure (PDB: 6FLN) (34,63). The alignment suggests that in the context of the full-length protein the RING domain comes in close proximity to the PRY/SPRY domain, explaining the importance of the interaction of CC and PRY/SPRY domain for TRIM25’s catalytic activity. Note the close proximity of the RING and the RNA binding site shown in blue. Catalytic activity of TRIM25, but not TRIM28, requires RING dimerization that in this model could only occur by association of two dimers. **(B).** A proposed potential mechanism of RNA-dependent activation of TRIM25 E3 ligase activity: The CC and PRY/SPRY of TRIM25 in the absence of RNA interact only transiently. Upon binding of stem-loop RNA, the PRY/SPRY domain and the CC interaction is stabilized. The lysine-rich linker also likely interacts with the stem (32). RIG-I after activation by viral dsRNA, dephosphorylation and possibly ubiquitination of the CTD by Riplet (2,22,23,82) is recruited to TRIM25 by an unknown mechanism, possibly involving binding the same RNA by both proteins (33). TRIM25 then poly-ubiquitinates K172 in the second CARD domain of RIG-I. In the absence of a suitable substrate, TRIM25 auto-ubiquitinates, possibly allowing for down-regulation of TRIM25 when not needed. This model assumes that distinct RNAs are necessary for association of TRIM25 and RIG-I and activation of RIG-I, as described for lnczc3h7a (33), but a single RNA might be sufficient for both. **(C)** A recent cryo-EM structure (PDB:7JLI) shows an interaction of the PRY/SPRY of a close relative of TRIM25, Riplet, with filaments of the helicase domain of RIG-I formed around double-stranded RNA (70).Overlay of this structure with the PRY/SPRY of TRIM25 shows, that the surface of Riplet equivalent to the one responsible for RNA binding in TRIM25 is facing away from the helicase domain and is not involved in contacting either RIG-I or dsRNA. It should however be noted that the RNA-binding residues of TRIM25 are not conserved in Riplet and is therefore possible that these otherwise closely related E3s have diverged evolutionary to bind RIG-I in an RNA-dependent way for TRIM25 and independent of RNA binding for Riplet.

We note that our proposed mechanism of TRIM25 activation resembles the mechanism of RNA dependent regulation of the E3-ubiquitin ligase activity of roquins (67): Roquins feature two rigid multidomain motifs, that are connected by a flexible linker and both bind RNA. RNA binding therefore removes flexibility from the system and forces the protein into an active conformation. It should be noted, that in this case the effect is E2 dependent, so that RNA binding not only changes the extent of ubiquitination, but also specificity for certain chain compositions. Similar effects might also occur for TRIM25, as the close proximity of the RNA binding site and the RING domain in the above model combined with the necessity to accommodate both RNA and the E2~ubiquitin conjugate likely cause steric restrictions for the full complex. Such an interference of the RING domain with RNA binding could explain the earlier observation that RNA binding of full-length TRIM25 is weaker than that of CC-PRY/SPRY alone (32).

We found no evidence for a direct interaction between the TRIM25 PRY/SPRY and RIG-I CARDs in our NMR experiments, although these two domains alone are sufficient for co-immunoprecipitation from cells (17). We noted however, that several of the mutants described to reduce the interaction between TRIM25 and RIG-I (F592, I594, L604 in murine TRIM25, corresponding to F597, I589 and L599 in human TRIM25) are located on or close to the second RNA binding site on the PRY/SPRY domain (53). This suggests that RNA binding of both proteins to double-stranded segments in the same RNA could play a crucial role in RIG-I activation (Figure 6B). This finding is supported by the recent identification of long non-coding RNAs that bind to both TRIM25 and RIG-I and facilitate their interaction (33,55). One of these RNAs is lnczc3h7a, that was described to interact with the RIG-I helicase domain independently of viral RNA and does not release autoinhibition of RIG-I (33).

Considering these new findings, we propose a mechanism, in which RNA-binding of TRIM25 not only assists in the recruitment of RIG-I through binding to the same RNA molecule but also directly activates the E3-ubiquitin ligase activity of TRIM25 by facilitating the interaction of the PRY/SPRY and CC domains (Figure 6B). Like RIG-I, many other substrates of TRIM25, e.g. ZAP, TUT4, MDM2, p53 and Dicer (29,30,61,68–70), are putative RNA binding proteins and a mechanism that involves recruitment of these substrates through binding to the same RNA molecule or enrichment in RNA granules and activation of the TRIM25 E3 ligase activity through RNA-binding might be more universal.

Several E3 ligases other than TRIM25 promote ubiquitination of RIG-I leading to controversial discussions about their relative importance (2,71–75). Among these are several TRIM proteins (TRIM4, 15, 40) and Riplet, a close relative of TRIM25, that lost the B-Box domains and parts of the CC (76–78). Mechanistically, a sequential ubiquitination of RIG-I by first Riplet in the CTD and then TRIM25 at the CARDs has been proposed (2). It is noteworthy that although Riplet is closely related to TRIM25, it has lost the regions on the CC and in the L2 linker, which harbours RNA binding in TRIM25, and the RNA binding lysines and histidine found in the PRY/SPRY are not conserved. A cryo-EM structure of the Riplet PRY/SPRY fused to filaments of the RIG-I helicase domain assembled around double-stranded RNA was recently published and gives insights into the likely interaction of RIG-I and Riplet (70). In this structure the regions of the Riplet-PRY/SPRY, that in TRIM25 are involved in RNA binding and harbour mutants that reduce RIG-I co-purification, are facing away from the helicase domain and play no role in the interaction (Figure 6C). Riplet-binding to RIG-I therefore seems to be independent of RNA. Together with earlier observations that Riplet preferentially ubiquitinates the CTD rather than the CARDs of RIG-I (73), this suggests that TRIM25 and Riplet interact with RIG-I in entirely different ways despite being closely related proteins and might have evolved to ubiquitinate RIG-I in an RNA dependent or independent manner respectively.

Interestingly, while the RNA-binding interface of TRIM25 is not conserved in Riplet at least one other E3 ligase ubiquitinating RIG-I, MEX3C, is an RBP (71), suggesting that a role of RNA-binding in substrate recruitment might be more common in other E3 ligases.

Despite the clear evidence for an activating role of RNA in RIG-I signalling, the role of Dengue virus subgenomic RNA in the inhibition of interferon expression remains unexplained (4). We found that TRIM25 binds a stem-loop of the Dengue virus subgenomic RNA with almost ten-fold higher affinity than the TRIM25-binding stem-loop of lnczc3h7a, a long non-coding RNA that binds both TRIM25 and RIG-I and thereby facilitates interferon expression (33). This suggests that the subgenomic RNA can outcompete lnczc3h7a for TRIM25 binding. It is however not clear what the effect of this competition is. RNA binding might also play a key role in other viral inhibition mechanisms for TRIM25. Most notably, the Influenza A non-structural protein 1 (NS1) also features an RNA-binding domain in addition to the TRIM25 binding effector domain (ED) (3,34). Interestingly, *in vitro* the ED alone is sufficient to explain binding to TRIM25, but in cells co-purification with TRIM25 depends on the presence of both domains (3,34). This suggests that binding to the same RNA could also stabilize the TRIM25/NS1 interaction, thus deactivating TRIM25-mediated ubiquitination. A similar mechanism was recently proposed for the interaction of NS1 with DHX30 (79). The inhibition of TRIM25 by other viral proteins such as paramyxovirus protein V and coronavirus protein N even depends only on their C-terminal domains that are also RBDs (5,8,80,81). RNA binding might therefore be widely exploited by viruses to inhibit TRIM25 and its crucial function in innate immunity.

## ACCESSION NUMBERS

Backbone assignments of the TRIM25-PRY/SPRY domain and extended PRY/SPRY domain have been deposited at the Biological Magnetic Resonance Bank (BMRB) under accession codes 27381 and 50226. SAXS data has been deposited in the SASBDB under accession codes SASDK78, SASDK88, SASDK98 and SASDKA8.

## ACKNOWLEDGEMENT

We thank the ESRF Grenoble (beamline BM29), EMBL/DESY Hamburg PETRA-III (P12 beamline), and the ILL (D22 beamline) local contacts for support (Taisiia Cheremnykh, Andrey Gruzinov, Al Kikhney, Gabriele Giachin and Anne Martel). This work was supported by the Francis Crick Institute which receives its core funding from Cancer Research UK (FC001142 to K.R.), the UK Medical Research Council (FC001142 to K.R.) and the Wellcome Trust (FC001142 to K.R.). J.H. gratefully acknowledges support via an Emmy-Noether Fellowship (HE 7291_1) and the Priority Program SPP1935 (EP37/3-1, 3-2) of the Deutsche Forschungsgemeinschaft (DFG) and the EMBL.

## FUNDING

This work was supported by the Francis Crick Institute which receives its core funding from Cancer Research UK (FC001142 to K.R.), the UK Medical Research Council (FC001142 to K.R.) and the Wellcome Trust (FC001142 to K.R.) and an Emmy-Noether Fellowship and a priority program to J.H., the Deutsche Forschungsgemeinschaft (DFG) [HE 7291_1, Priority Program SPP1935 to J.H.]).

## CONFLICT OF INTEREST

The authors declare no competing interests.

## Notes

### Competing Interest Statement

The authors have declared no competing interest.

