## Supplementary Data for "Mechanistic insights into RNA binding and RNA-regulated RIG-I ubiquitination by TRIM25"

Supplementary Table 1. Small-angle X-ray scattering statistics

| | **TRIM25 CC-PRY/SPRY** | **TRIM25 CC-PRY/SPRY/pre-let-7** | **TRIM25 CC-PRY/SPRY/ lnczc3h7a-SL** | **TRIM25 CC-PRY/SPRY/pre-let-7** |
|---|---|---|---|---|
| (a) Sample Details | | | | |
| Organism | *Homo sapiens* | | | |
| Source | *E. coli* BL2 (DE3) | *E. coli* BL2 (DE3)/*in vitro* transcription | | |
| Uniprot sequence ID | Q14258 | | | |
| Description | TRIM25 189-630 | TRIM25 189-630 in complex with the pre-let-7 stem-loop | TRIM25 189-630 in complex with the lnczc3h7a stem-loop | TRIM25 189-630 in complex with the pre-let-7 stem-loop |

| | | | | |
|---|---|---|---|---|
| Molecular mass M from chemical composition (Da) | 49,952 | 58,195 | 57,358 | 58,195 |
| loading concentration (mg/ml) | 0.37, 0.75, 1.5, 3 and 6 | n.d. | n.d. | n.d. |
| injection volume (µl) | 30 | 100 | 30 | 500 |
| concentration (µM) | 7.4, 15, 30, 60, 120 | n.d. | n.d. | n.d. |
| Solvent composition and source | 20 mM MES, pH 6.5, 75 mM NaCl and 0.5 mM TCEP | | | |

(b) SAS data collection parameter

| | | | | |
|---|---|---|---|---|
| Source and instrument | Hamburg PETRA-III P12 with Dectris Pilatus 6M | | | ESRF BM29 with Dectris Pilatus |
| Wavelength (Å) | 1.22 | 1.24 | 1.24 | 0.99 |
| Sample-detector distance (m) | 3.0 | 3.0 | 3.0 | 2.869 |
| q-measurement range (Å) | 0.0226-7.405 | 0.0252-7.318 | 0.0297-7.267 | 0.0346-4.928 |
| Radiation damage monitoring | frame-by-frame comparison | | | |
| Exposure time (s) & number | 0.05x20 | 0.195x100 | 0.05x20 | 1s |

| | | | | |
|---|---|---|---|---|
| Sample configuration | sample changer with flow through capillary measurement | | | SEC-SAXS, Superdex S200 10/300 Increase |
| Sample temperature (℃) | 20 | 25 | 20 | 20 |

(c) Software employed for SAS data reduction, analysis and interpretation

| | | | | |
|---|---|---|---|---|
| SAXS data processing | I(q) vs. q using Bsx cube, solvent subtraction and curve merging using PRIMUSqt from ATSAS (Franke et al., 2017) | | | I(q) vs. q using Bsx cube, solvent subtraction and curve merging Chromixs |
| Basic analyses: Guinier, P(r), Vp | PRIMUSqt from ATSAS 2.7.1 (Franke et al., 2017) | | | |
| Atomic structure modelling | CRYSOL 2.8.2 from PRIMUSqt in ATSAS 2.8 (Svergun et al., 1995) | | | |
| Molecular graphics | -- | -- | - | - |

(d) Structural parameters

| | | | | |
|---|---|---|---|---|
| | Guinier analysis | | | |
| I(0) (raw) | 51290 ± 160 | 34043 ± 42 | 0.033 ± 0.001 | 21.84 ± 0.22 |
| $R_g$ (Å) | 68.3±0.5 | 56.3 ± 0.1 | 58.1 ± 1.8 | 57.0 ± 0.5 |
| $qR_g$ max | 1.50 | 1.4 | 1.28 | |
| Coefficient of correlation, $R^2$ | 0.82 | 0.80 | 0.77 | 0.77 |
| | P(r) Analysis from AUTOGNOM | | | |

| | | | | |
|---|---|---|---|---|
| I(0) (cm-1) | 54320 | 35340 | 0.034 | 243 |
| $R_g$ (Å) | 78.0 | 61.3 | 56.4 | 66.0 |
| $d_{max}$ (Å) | 305.6 | 226.6 | 165.6 | 205.0 |
| *q* range ($Å^{-1}$) | 0.202-3.06 | 0.148-2.50 | 0.145-1.38 | 0.394-3.31 |
| $\chi^2$ (total estimate from *GNOM*) | 0.60 | 0.73 | 0.68 | 0.60 |
| Porod volume ($Å^{-3}$) (ratio $V_P$/calculated *M)* | 302540 | 182720 | 180000 | 215000 |

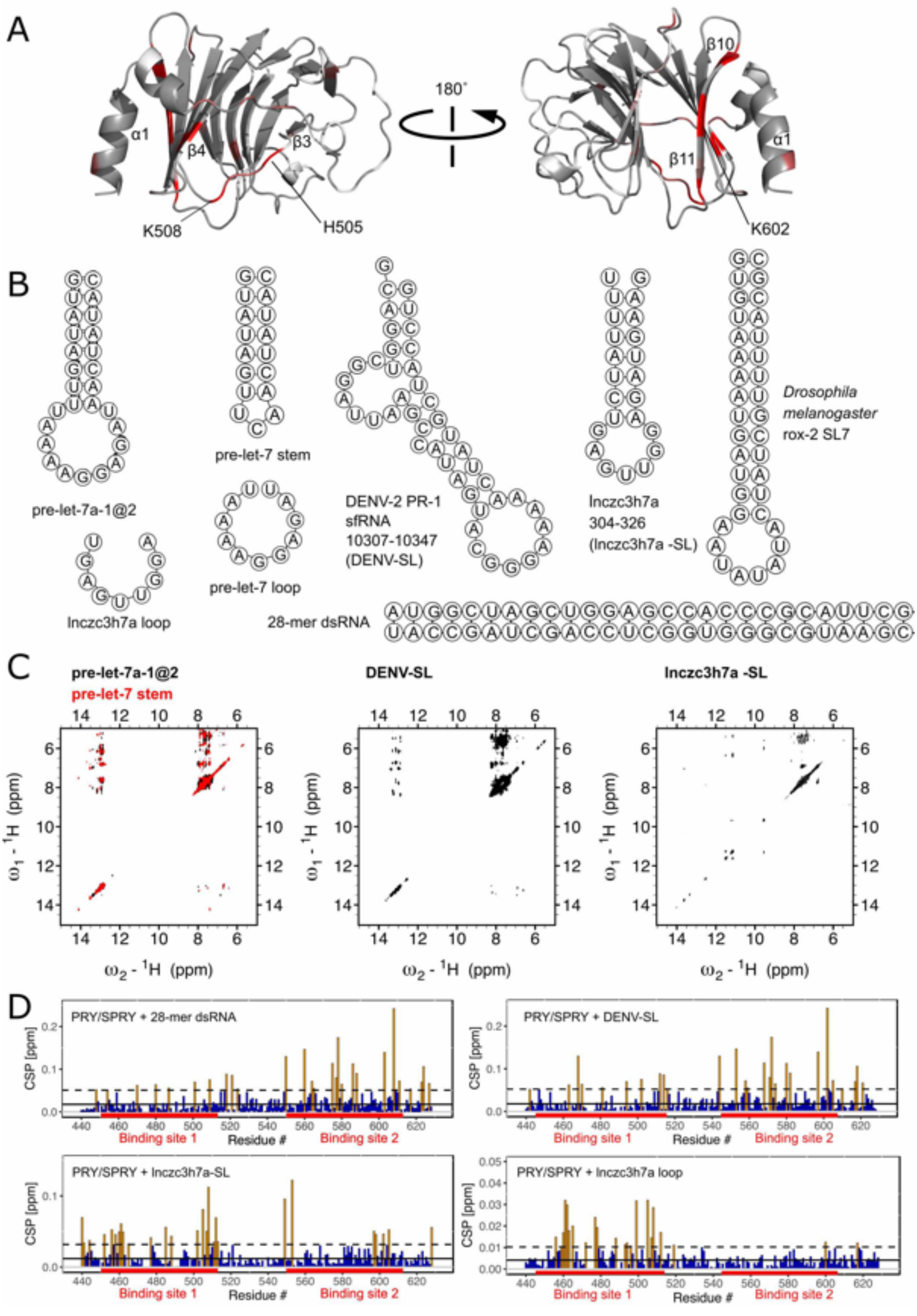
A
α1
β4
β3
K508
H505
180°
β10
β11
α1
K602
B
pre-let-7a-1@2
pre-let-7 stem
pre-let-7 loop
DENV-2 PR-1
sfRNA
10307-10347
(DENV-SL)
lnczc3h7a
304-326
(lnczc3h7a -SL)
Drosophila
melanogaster
rox-2 SL7
lnczc3h7a loop
28-mer dsRNA
C
pre-let-7a-1@2
pre-let-7 stem
DENV-SL
lnczc3h7a -SL
$\omega_1$ - $^1$H (ppm)
$\omega_2$ - $^1$H (ppm)
D
PRY/SPRY + 28-mer dsRNA
PRY/SPRY + DENV-SL
PRY/SPRY + lnczc3h7a-SL
PRY/SPRY + lnczc3h7a loop
CSP [ppm]
Residue #
Binding site 1
Binding site 2

Supplementary Figure S1: Structure of RNAs used in this study and their interaction with TRIM25 PRY/SPRY. **(A)** NMR titrations show two separate RNA binding sites on the PRY/SPRY: CSPs from titration of TRIM25 PRY/SPRY with pre-let-7 plotted on the structure (PDB: 6FLM) indicate two binding sites. Binding-site-1 is formed by ß-strand 4 and the flexible linker connecting it with ß-strands 3. Binding-site-2 is formed by ß-strands 10 and 11. (**B**) Predicted structures of the RNAs used in this study. All RNA structures were predicted using the RNAfold webserver (75). **(C)** The observation of peaks in the imino region (10-15 ppm) in $^{1}H/^{1}H$-2D-NOESYs confirms that the stem-loops indeed form double-stranded regions. Overlay of the NOESYs of pre-let-7a-1@2 and the truncated pre-let-7 stem show very similar imino-regions indicating that both constructs form similar stems **(D)** NMR titrations with RNAs of different structure: Titration with a 28-mer duplex RNA shows in addition to strong signal loss CSPs mostly affecting binding site 2, supporting a specificity of this binding site for double-stranded RNA. The stem-loops found in the subgenomic RNA of the Dengue virus and the long-noncoding RNA lnczc3h7a shows additional CSPs in binding site 1. The loop region of lnczc3h7a binds selectively to binding site 1 on the PRY/SPRY, similar to the loop of pre-let-7, but with weaker affinity.

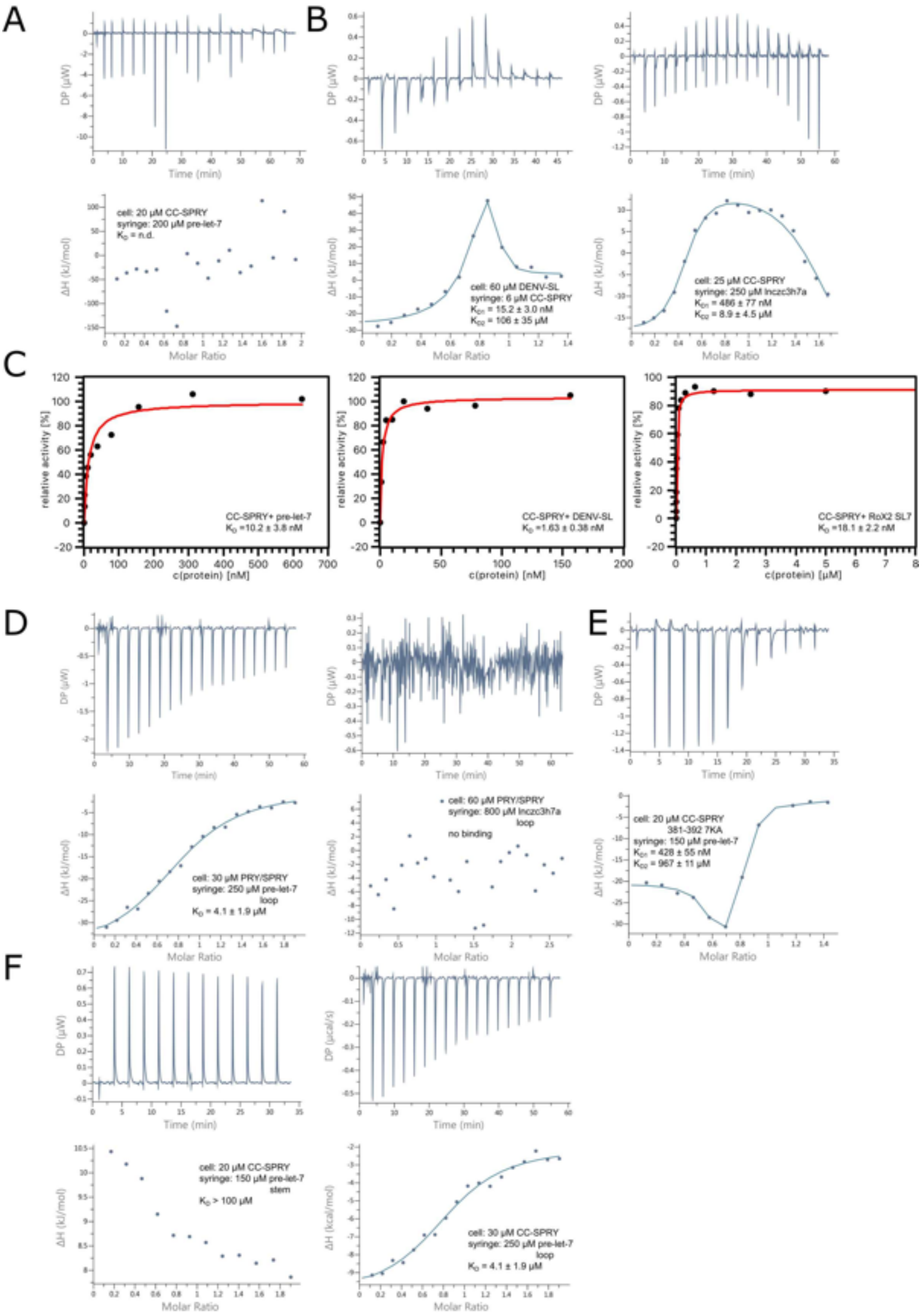
A
B
cell: 20 µM CC-SPRY
syringe: 200 µM pre-let-7
K_D = n.d.
cell: 60 µM DENV-SL
syringe: 6 µM CC-SPRY
K_D1 = 15.2 ± 3.0 nM
K_D2 = 106 ± 35 µM
cell: 25 µM CC-SPRY
syringe: 250 µM lnczc3h7a
K_D1 = 486 ± 77 nM
K_D2 = 8.9 ± 4.5 µM
C
CC-SPRY+ pre-let-7
K_D =10.2 ± 3.8 nM
CC-SPRY+ DENV-SL
K_D =1.63 ± 0.38 nM
CC-SPRY+ RoX2 SL7
K_D =18.1 ± 2.2 nM
relative activity [%]
c(protein) [nM]
c(protein) [µM]
D
E
cell: 30 µM PRY/SPRY
syringe: 250 µM pre-let-7 loop
K_D = 4.1 ± 1.9 µM
cell: 60 µM PRY/SPRY
syringe: 800 µM lnczc3h7a loop
no binding
cell: 20 µM CC-SPRY 381-392 7KA
syringe: 150 µM pre-let-7
K_D1 = 428 ± 55 nM
K_D2 = 967 ± 11 µM
F
cell: 20 µM CC-SPRY
syringe: 150 µM pre-let-7 stem
K_D > 100 µM
cell: 30 µM CC-SPRY
syringe: 250 µM pre-let-7 loop
K_D = 4.1 ± 1.9 µM
DP (µW)
DP (µcal/s)
Time (min)
ΔH (kJ/mol)
ΔH (kcal/mol)
Molar Ratio

Supplementary Figure S2: TRIM25/RNA interactions quantified by ITC and filter binding assays **(A)** ITC with TRIM25 CC-PRY/SPRY and pre-let-7 shows complex binding modes for the stem-loops with two or more binding sites **(B)** ITC of CC-PRY/SPRY titrated to DENV-SL (left panel) and lnczc3h7a titrated to CC-PRY/SPRY (right panel). **(C)** Filter-binding assays confirm binding of TRIM25 CC-PRY/SPRY with low nanomolar affinity to pre-let-7 and DENV-SL. In addition, they also show binding to roX2 SL7, a functionally unrelated non-coding RNA from *Drosophila melanogaster,* that shows little sequence similarity, but forms a similar stem-loop, suggesting that RNA structure might be more important for specificity than sequence. **(D)** ITC data for TRIM25 PRY/SPRY titrated by pre-let-7 loop (left panel) and lnczc3h7a loop (right panel). **(E)** ITC of CC-PRY/SPRY 381-392 7KA titrated by pre-let-7. **(F)** ITC of CC-PRY/SPRY titrated by pre-let-7 stem (left) and loop (right).

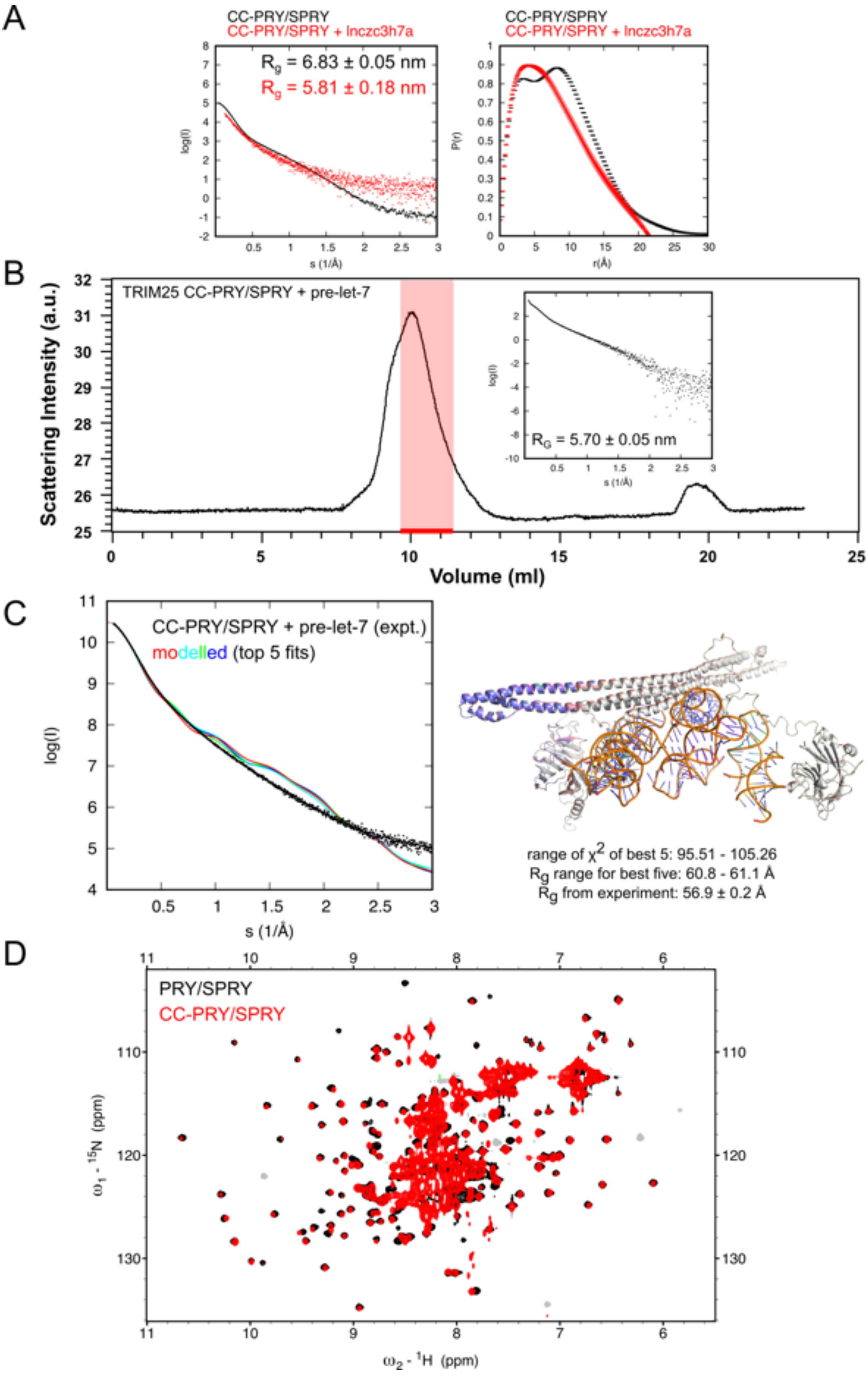
A
CC-PRY/SPRY
CC-PRY/SPRY + lnczc3h7a
Rg = 6.83 ± 0.05 nm
Rg = 5.81 ± 0.18 nm
log(I)
s (1/Å)
P(r)
r(Å)
B
TRIM25 CC-PRY/SPRY + pre-let-7
Scattering Intensity (a.u.)
Volume (ml)
RG = 5.70 ± 0.05 nm
C
CC-PRY/SPRY + pre-let-7 (expt.)
modelled (top 5 fits)
range of χ2 of best 5: 95.51 - 105.26
Rg range for best five: 60.8 - 61.1 Å
Rg from experiment: 56.9 ± 0.2 Å
D
PRY/SPRY
CC-PRY/SPRY
ω1 - 15N (ppm)
ω2 - 1H (ppm)

Supplementary figure S3: CC and PRY/SPRY cooperate to bind RNA. **(A)** Binding of lnczc3h7a-SL leads to a similar reduction of the radius of gyration and similar changes to the pair-wise distribution function in SAXS of CC-SPRY/SPRY than pre-let-7, indicating that stem-loop binding generally leads to the formation of a more compact complex. **(B)** This is also confirmed by SEC-SAXS data showing that this reduction is not due to the presence of free RNA. **(C)** Molecular modelling starting from a structure that fits well to the SAXS curve of free TRIM25 CC-PRY/SPRY (first described in Koliopolous et al. 2018) shows that addition of RNA alone, even when happening close to the center of mass at the middle of the CC cannot explain the observed reduction in $R_g$. **(D)** Overlay of the $^{1}H/^{15}N$-HSQCs of TRIM25 PRY/SPRY and CC-PRY/SPRY show excellent overlap for the dispersed peaks corresponding to folded regions. Signal intensity for these peaks is however weaker at the same concentration for the CC-PRY/SPRY than it is for the PRY/SPRY. Together with the lack of CSPs this suggests that only the unbound state of the PRY/SPRY is observed while the state of PRY/SPRY bound to the CC is not.

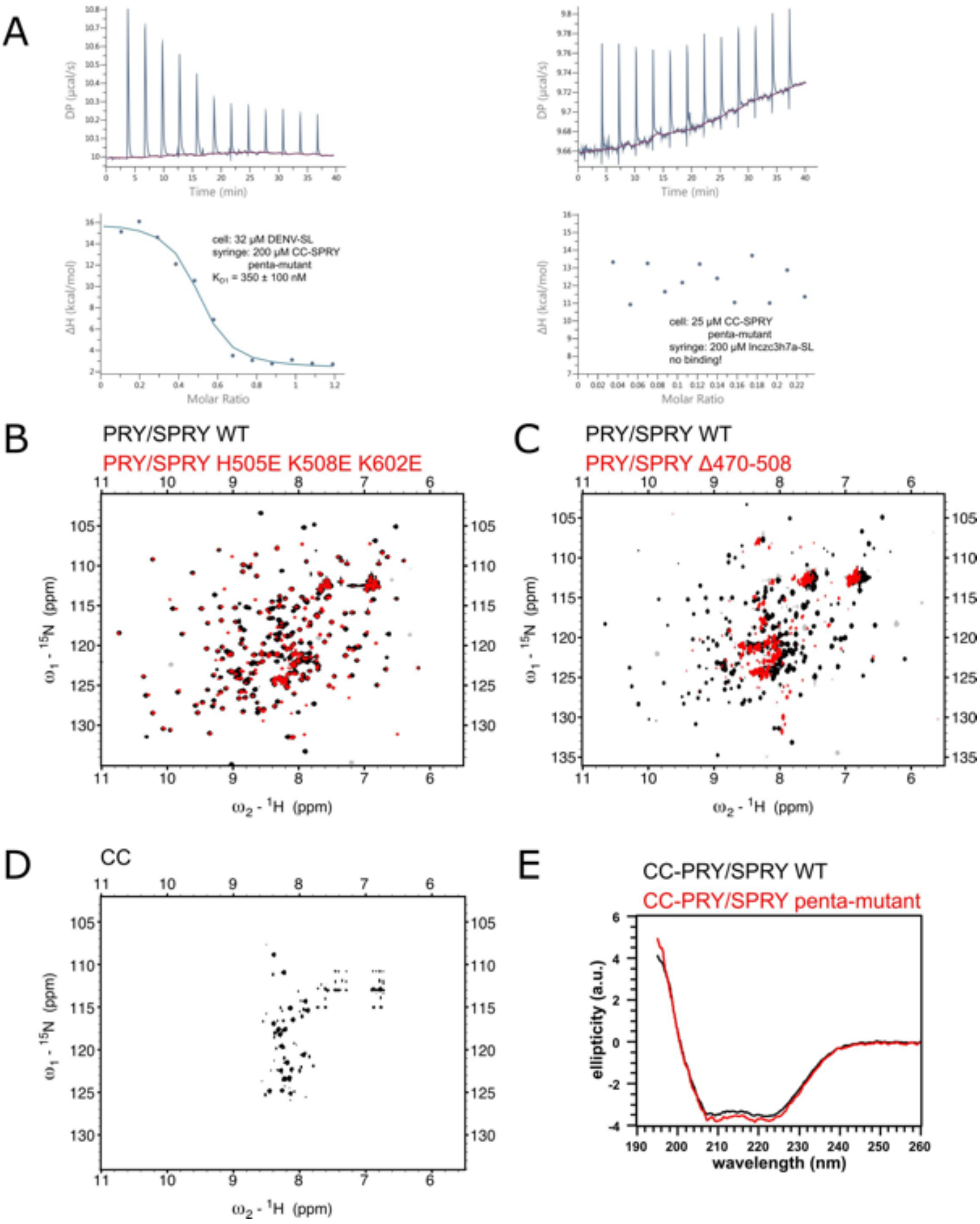


Supplementary figure S4: Mutation of the RNA binding interface. **(A)** ITC shows that the penta-mutant disrupts also the binding of DENV-SL and lnczc3h7a-SL. **(B)** The $^1$H/$^{15}$N-HSQC of the triple mutant on the PRY/SPRY described here is very similar to the wildtype with only few peaks significantly shifted indicating only minor local perturbations to the structure. (C) In contrast the $^1$H/$^{15}$N-HSQC of the previously described Δ470-508 deletion in the PRY/SPRY domain (ΔRBD by Choudhury et al. 2017)

lacks the well dispersed peaks of a well-folded, β-strand rich protein, indicating that the deletion of this region disrupts folding of the PRY/SPRY domain. (D) **)** The $^{1}H/^{15}N$-HSQC of the CC show only peaks corresponding to the flexible termini due to its large size and elongated shapee, which leads to fast transverse relaxation of structured regions. Therefore, NMR is not suitable to assess the structural impact of mutants on the CC. (E) The circular dichroism spectra of the penta-mutant also overlay well with the wild type suggesting that the mutants on both CC and PRY/SPRY do not adversely affect protein folding.

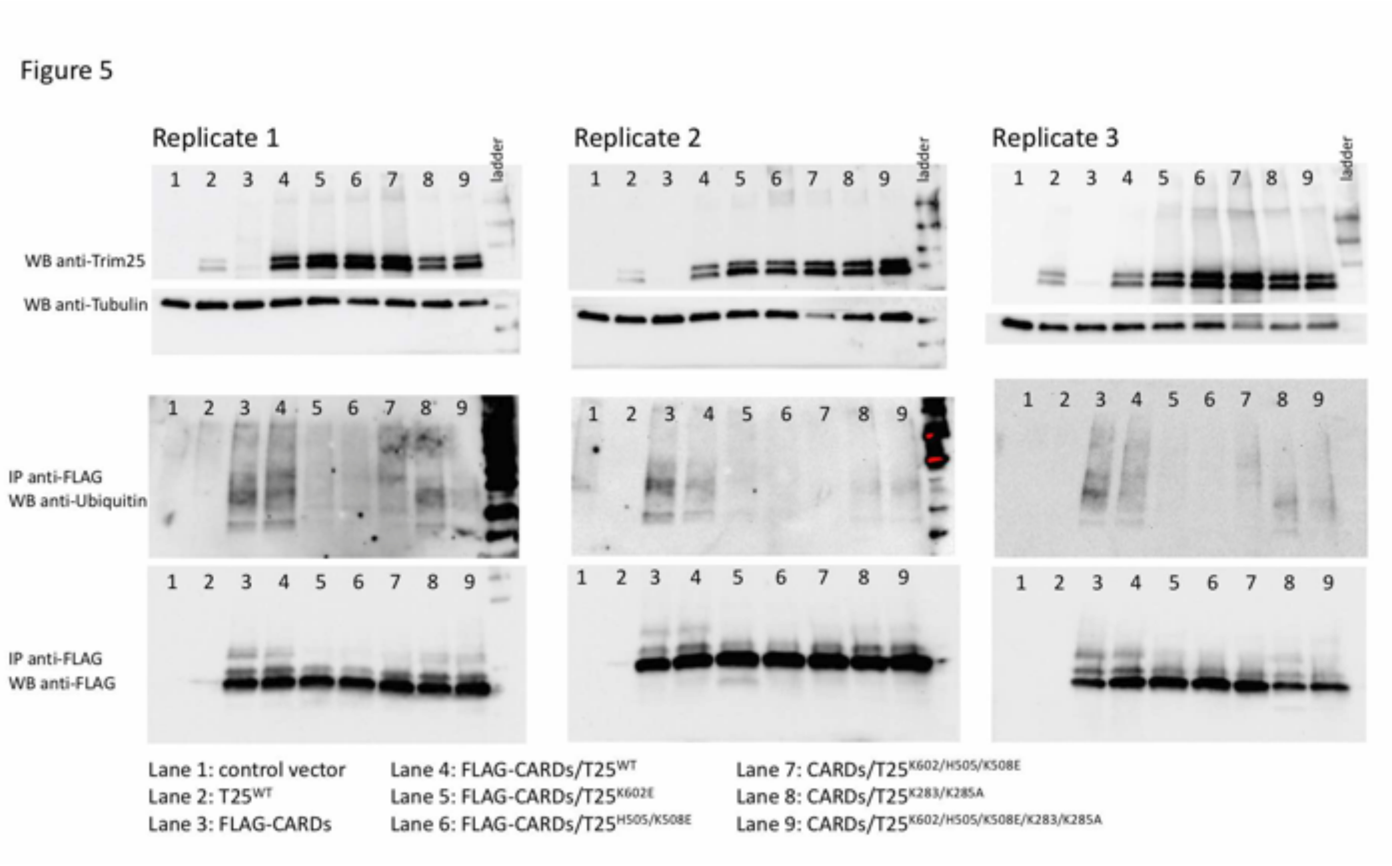


Supplementary Figure 5: Replicate Western bots for Ubiquitination assays from Figure 4.
